# Dynamics of genomic change during evolutionary rescue in the seed beetle *Callosobruchus maculatus*

**DOI:** 10.1101/364158

**Authors:** Alexandre Rêgo, Frank J. Messina, Zachariah Gompert

## Abstract

Rapid adaptation can prevent extinction when populations are exposed to extremely marginal or stressful environments. Factors that affect the likelihood of evolutionary rescue from extinction have been identified, but much less is known about the evolutionary dynamics (e.g., rates and patterns of allele frequency change) and genomic basis of successful rescue, particularly in multicellular organisms. We conducted an evolve-and-resequence experiment to investigate the dynamics of evolutionary rescue at the genetic level in the cowpea seed beetle, *Callosobruchus maculatus*, when it is experimentally shifted to a stressful host plant, lentil. Low survival (∼1%) at the onset of the experiment caused population decline. But adaptive evolution quickly rescued the population, with survival rates climbing to 69% by the F5 generation and 90% by the F10 generation. Population genomic data showed that rescue likely was caused by rapid evolutionary change at multiple loci, with many alleles fixing or nearly fixing within five generations of selection on lentil. Selection on these loci was only moderately consistent in time, but parallel evolutionary changes were evident in sublines formed after the lentil line had passed through a bottleneck. By comparing estimates of selection and genomic change on lentil across five independent *C. maculatus* lines (the new lentil-adapted line, three long-established lines, and one case of failed evolutionary rescue), we found that adaptation on lentil occurred via somewhat idiosyncratic evolutionary changes. Overall, our results suggest that evolutionary rescue in this system can be caused by very strong selection on multiple loci driving rapid and pronounced genomic change.

## Introduction

Decades of field and lab studies have shown that adaptation can be rapid, in some cases occurring over a few to several generations (e.g., Steinhauer & Holland, 1987; Grant & Grant, 2002; Thompson, 2013; Bergland *et al*., 2014; Elmer *et al*., 2014; Nosil *et al*., 2018). Evidence for rapid adaptive evolution is particularly common in human-altered environments (e.g., during adaptation to pesticides, antibiotics, or pollution; Palumbi, 2001; Vonlanthen *et al*., 2012; Cook & Saccheri, 2013) or when adaptation is driven by interactions among species (e.g., resource competition, host-pathogen interactions, or predator-prey interactions; Yoshida *et al*., 2003; Stuart *et al*., 2014; Antonio-Nkondjio *et al*., 2015; Behrman *et al*., 2018). Rapid adaptive evolution may also prevent sustained demographic decline and extinction when populations are exposed to extremely marginal or stressful environments during a process known as evolutionary rescue (Gomulkiewicz & Holt, 1995; Bell & Gonzalez, 2009; Gonzalez *et al*., 2013; Lindsey *et al*., 2013; Orr & Unckless, 2014). Whereas most theory and experiments have focused on the probability of evolutionary rescue under different conditions (reviewed in Bell, 2017), much less is known about the evolutionary dynamics (e.g., rates and patterns of evolutionary change over time at individual loci or across the genome) and genomic consequences of rescue when it occurs (but see Wilson *et al*., 2017).

Evolutionary rescue differs from other forms of adaptive evolution in a few key ways that could result in distinct evolutionary dynamics and genomic signals. First, evolutionary rescue couples ecological and evolutionary dynamics, because low absolute fitness in a deteriorating or stressful environment causes population decline that is then reversed when evolution leads to a sufficiently large increase in absolute fitness (Gomulkiewicz & Holt, 1995; Orr & Unckless, 2014). Second, compared to other cases of adaptive evolution, evolutionary rescue necessarily occurs in populations far from a phenotypic optimum (because population decline implies a poor fit to the current environment). Thus, mutations of major effect could contribute disproportionately to evolutionary rescue (McKenzie & Batterham, 1994; Orr, 2005). Consistent with this prediction, theory (e.g., Fisher’s geometric model; Fisher, 1938; Orr, 1998) and experimental evolution studies (e.g., Lenski *et al*., 1991; Barrick *et al*., 2009) suggest that mutations of major effect drive early stages of adaptation. Recent theory also suggests that evolutionary rescue is more likely when standing genetic variation is present, and may often involve soft selective sweeps in which multiple beneficial mutations increase in frequency simultaneously (Hermisson & Pennings, 2005; Bell, 2017; Wilson *et al*., 2017). Thus, substantial genetic variation might be retained in a population throughout this process.

Because evolutionary rescue often involves rapid adaptation (e.g., Bell & Gonzalez, 2009; Bell, 2013; Vander Wal *et al*., 2013; Kreiner *et al*., 2017), cases of rescue could provide tractable opportunities to study the dynamics of adaptive alleles during a complete bout of adaptation, that is, from the onset of population decline to when a population has rebounded demographically. Such studies should also help determine whether instances of repeated ecological dynamics (e.g., population decline and recovery) are driven by repeatable evolutionary dynamics, and thus whether eco-evolutionary dynamics are repeatable or predictable (Rudman *et al*., 2018). Whereas experimental studies have documented patterns of ecological and evolutionary change during rescue (e.g., Bell & Gonzalez, 2009; Gonzalez & Bell, 2013; Ramsayer *et al*., 2013; Killeen *et al*., 2017), such work has mostly focused on microorganisms (but see, e.g., Agashe, 2009; Agashe *et al*., 2011) and has rarely been combined with genetic or genomic data. Here, we conduct an evolve-and-resequence experiment to investigate the dynamics and outcome of evolutionary rescue at the genetic level in the cowpea seed beetle, *Callosobruchus maculatus* (Chrysomelidae), when it is experimentally shifted to a marginal host plant, lentil (*Lens culinaris*, Fabaceae).

*Callosobruchus* beetles infest human stores of grain legumes. Females attach eggs to the surface of legume seeds. Upon hatching, larvae burrow into and develop within a single seed. Because *C. maculatus* has been associated with stored legumes for thousands of years, laboratory conditions are a good approximation of its “natural” environment (Tuda *et al*., 2014). Beetle populations mainly attack grain legumes in the tribe Phaseoleae, particularly those in the genus *Vigna* (Tuda *et al*., 2006). Lentil (*L. culinaris*), a member of the tribe Fabeae, is a poor host for most *C. maculatus* populations, as larval survival in seeds is typically <5% (Messina *et al*., 2009a,b, 2018). However, lentil is used as a host by a few unusual ecotypes (Credland, 1987, 1990). Previous attempts to establish laboratory populations on lentil have often resulted in extinction (Credland, 1987), but in a few cases experimental lines have rapidly adapted to lentil (Messina *et al*., 2009b). For example, in three experimental lines, survival rose from ∼2% to >80% within 20 generations, and these lines have now persisted on lentil for >100 generations (Gompert & Messina, 2016). Thus, evolutionary rescue can occur in this system.

In the current study, we established a new lentil-adapted line (denoted L14), which we then split into two sublines (L14A and L14B) before evolutionary rescue was complete, that is, after the population began to rebound from an initial bottleneck, but before it reached a performance plateau (Fig. 1). We sampled and sequenced beetles nearly every generation, and could thus characterize genome-wide evolutionary dynamics on a fine temporal scale. Our goal was not to identify specific genes that mediate evolutionary rescue, but rather to determine (i) whether rescue depends on a few or many genetic loci, (ii) whether selection on individual genetic loci is consistent throughout the process (across time and between sublines), and (iii) whether selection causes alleles to fix or instead causes more subtle shifts in allele frequencies (i.e., partial or incomplete sweeps), as has been observed during other evolve-and-resequence experiments with multicellular organisms (e.g., Burke *et al*., 2010; Graves Jr *et al*., 2017). We focus on a single lentil line, because other recent attempts to establish lentil lines failed (>10 lines started between October 2013 and September 2014), resulting in extinction. This precludes us from directly asking about the repeatability of genomic change during evolutionary rescue and of using parallel change as evidence of selection (we instead rely on models that account for the effects of drift when inferring selection). However, we are able to compare patterns of genomic change during rescue for this single new lentil line with the genomic outcomes of rescue in three independently derived, long-established lentil lines (lines L1, L2 and L3), and with a case of failed evolutionary rescue from one of our other recent attempts to establish *C. maculatus* on lentil (hereafter, line L11) (Fig. 1a). We treat these as exploratory comparisons designed to provide preliminary insights into the ways in which patterns of genome-wide evolutionary change in *C. maculatus* on lentil might be repeatable.

**Figure 1:**
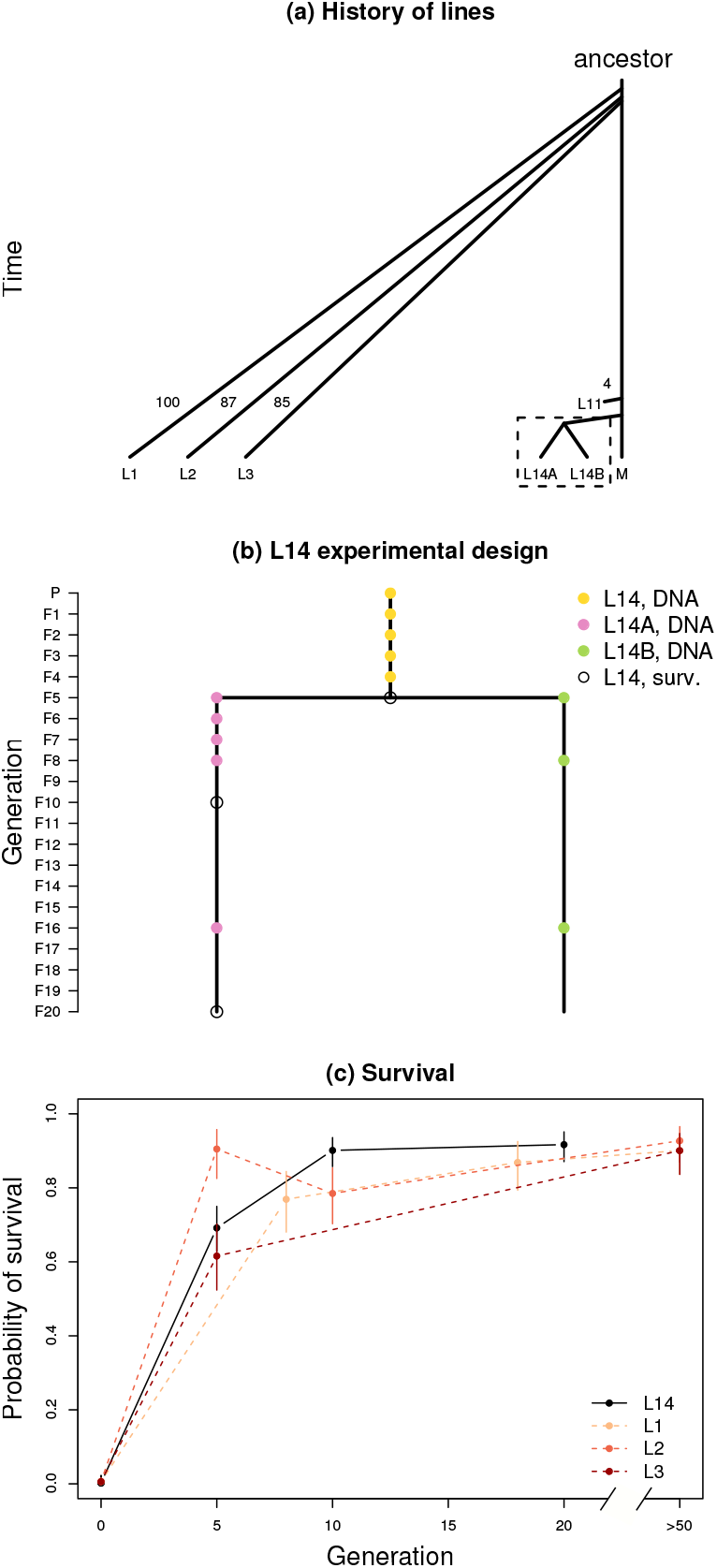
Experimental design. Panel (a) illustrates the history of all lentil lines discussed in this manuscript (i.e., L1, L2, L3, L11 and L14) along with the south Indian mung line (denoted M). The number of generations that elapsed between the origin of each line and when it was sampled for population genomic analyses is shown for L1, L2, L3 and L11. Details for L14 (denoted by the box) are shown in panel (b). The L14 lentil line was established from an Indian mung bean line. At the F5 generation, L14 was split into sublines A and B. Samples were taken for genetic analysis every generation up to F4 (yellow dots), and then in subline A in the F5–F8, and F16 generations (pink dots), and subline B in the F5, F8, and F16 generations (green dots). Open black circles denote generations in which fitness was assayed. Bayesian estimates of survival on lentil are shown in panel (c). Survival was measured at generations L14–F5, L14A–F10, L14A–F20, and in the Indian mung bean line, which is shown as generation 0. Data for L1, L2, and L3 come from Messina *et al*. (2009b) and Messina & Durham (2015). Points and vertical lines denote posterior medians and 95% equal-tail probability intervals.

## Materials & Methods

### Study system, selection experiment and fitness assays

The new line (L14) produced for the current study, the three long-established lentil lines (L1-L3) (∼100 generations on lentil), and the line that failed to adapt to lentil (L11) were derived from the same base population of *C. maculatus* that was originally collected from southern India (Messina, 1991; Mitchell, 1991) (Fig. 1a). This base population had been continuously reared on mung bean, *Vigna radiata* (L.) Wilczek, for >300 generations at the time we began this experiment. Previous assays demonstrated that, for this Indian beetle population, initial survival to adult emergence is only 1-2% in lentil (Messina *et al*., 2009b; Messina & Jones, 2011). Consequently, there is always a severe initial bottleneck, and most attempts to produce a self-sustaining population on lentil seeds eventually fail (Messina *et al*., 2009a; Gompert & Messina, 2016). In the lines designated L1-L3, survival increased rapidly over the course of only a few generations. Survival reached >60% after only five generations, and >80% in fewer than 20 generations (Messina *et al*., 2009b). At the same time, there were substantial reductions in development time and increases in body size. Genomic analyses of these lines, which focused on genetic trade-offs for adaptation to mung bean versus lentil, did not commence until each had been maintained on lentil for 80-100 generations, and had reached a plateau with respect to performance on the novel host (Gompert & Messina, 2016). Hence, we did not capture the initial stages of adaptation in the earlier study. This can be important, as early stages of rescue can have an sizeable effect on outcomes (e.g., Lagator *et al*., 2014).

The three lentil-adapted lines (L1-L3) were established as described by Messina *et al*. (2009a,b). We followed the same protocol in the current study. Our goal was to establish one or more lentil-adapted lines, as this would allow us to sample the line(s) every generation for population genomic analyses during the early phase of adaptation (as described below). As expected, multiple (>10) initial attempts to produce a new lentil-adapted line eventually resulted in population extinction (including L11), but a single line (L14) exhibited the rapid rise in survival previously observed in L1-L3 (see Results). This line was formed by adding >4000 founding adults to 1500 g of lentil seeds (about 24,000 seeds). Most F1 offspring emerged 55–65 days after the founding adults were added. We transferred F1 beetles (approximately 100–200 individuals) to a new jar to form the F2 generation. Following the severe bottleneck in the initial generation on lentil, larval survival in seeds increased rapidly (as described below), so that we were able to use at least a few hundred beetles to form each successive generation (as in past work, transfers were made during the peak of emergence to avoid artificial selection for rapid or delayed development; Messina *et al*., 2009b). After five generations, the L14 population size was sufficiently high to implement standard culturing techniques, which involved transferring >2000 beetles to a new batch of 750g lentil seeds each generation (see “Culturing and establishing lines” in the OSM). At the F5 generation, the L14 line was split into sublines A and B (Fig. 1b) (this was the first generation where a sufficient number of beetles emerged to subdivide the line). By doing so, we could assess whether the evolutionary dynamics after a shared bottleneck were repeatable (i.e., parallel). While our experiment achieved some replication by using two sublines and by conducting comparisons with the older lentil lines and one of the failed new lines, we lack replication for evolutionary dynamics during the early stages of (successful) adaptation (i.e., of evolutionary rescue). Nonetheless, even one or a few instances of adaptation can provide important insights into how evolution can occur (examples of this include the evolution of beak size in response to drought in Darwin’s finches and the evolution of citrate metabolism in *E. coli*, with the latter occurring in only one of a dozen replicate lines; Grant & Grant, 2002; Blount *et al*., 2008).

By the F5 generation, the population size of the L14 line was sufficiently high to apply our standard protocol for measuring survival in lentil from egg hatch to adult emergence (Messina *et al*., 2009b). We established a cohort of larvae in lentil seeds by first placing three pairs of newly emerged adults into each of 40 petri dishes containing about 100 lentil seeds. After 10-15 days, we collected a few seeds bearing a single hatched egg from each dish, and isolated each seed in a 4-ml vial. Vials were inspected daily for adult emergence until two weeks after the last adult had emerged. We collected a total of 224, 224, and 182 infested seeds for assays of the F5, F10, and F20 generations (Fig. 1b). To reduce any effects of parental host, the L14 line was reverted back to mung bean for one generation (i.e., parents of all test larvae had developed in mung bean) (Messina *et al*., 2009a). Survival probabilities were estimated using a Bayesian binomial model with an uninformative (Jefferys) beta prior on the survival proportions. This model has an analytical solution, *Pr*(*p|y, n*) = beta(*α* = *y* + 0.5*, β* = *n* − *y* + 0.5), where *p* is the survival probability and *y* is the number of beetles that survived out of the *n* infested seeds. Thus, exact posteriors are presented.

### Genetic data

We sampled and isolated genomic DNA from 48 adult beetles per generation for the L14 founders (the P generation) as well as for the F1-F4 generations. After L14 line was split into two sublines (A and B) we sampled beetles from subline A (L14A) at generations F5, F6, F7, F8 and F16, and from subline B (L14B) at generations F5, F8 and F16 (Fig. 1b). We also sampled and isolated DNA from the failed F11 line; this was done in the F4 generation only (48 individuals were sampled; see “The L11 line” in the OSM for additional details). We generated partial genome sequences for these 672 *C. maculatus* beetles using our standard genotyping-by-sequencing approach (see “Our GBS approach” in the OSM; Gompert *et al*., 2012, 2014b). This approach provides a sample of SNPs distributed across the genome. We do not assume that the actual alleles responsible for lentil adaptation are included in this set of SNPs, but we do expect these data to include SNPs indirectly affected by selection on the causal genetic loci through linkage disequilibrium (LD). This genomic sampling scheme should thus provide a reasonable approximation of the evolutionary dynamics of causal variants. Our approach differs in some respects from most evolve-and-resequence experiments that used pooled whole genome sequencing of samples taken at a few time points (e.g., Burke *et al*., 2010; Orozco-terWengel *et al*., 2012; Tobler *et al*., 2014; Graves Jr *et al*., 2017). By foregoing the expenses associated with whole-genome sequencing (the standard approach), we were able to obtain (partial) genome sequence data that were tied to individual seed beetles, and we were able to sample nearly every generation during adaptation. These individual-level data were critical for confidently measuring LD among loci. Moreover, because adaptation was so rapid, we likely would have missed most of the dynamics of adaptation without our fine-scale temporal sampling (see Results and Discussion). The might not be a problem in systems where adaptation occurs more slowly, but it is hard to know the pace of adaptation without such temporal resolution.

We used the `aln` and `samse` algorithms from `bwa` (ver. 0.7.10) (Li & Durbin, 2009) to align the 764 million ∼86 bp DNA sequences (after trimming barcodes) to a new draft genome assembly for *C. maculatus* (Fig. S1; see “*De novo* assembly of a *C. maculatus* genome” and “Alignment and variant calling” in the OSM for details). We then identified SNPs using the Bayesian multiallelic/rare variant caller from `samtools` (version 1.5) and `bcftools` (version 1.6) (implemented with the `-m` option in `bcftools call`). SNPs were subsequently filtered based on a variety of criteria, including minimum mean coverage (≈2*⇥* per beetle), maximum percent missing data (30%), and mapping quality (30) (see the OSM for details). We retained 21,342 high-quality SNPs after filtering (about 1 SNP per 50 kpbs). Genetic data from the long-established lentil lines (L1, L2, and L3) were described in Gompert & Messina (2016). These samples were collected after 100 (L1), 87 (L2) and 85 (L3) generations of evolution on lentil (N = 40 individuals per line), and also include a reference sample from the source mung bean line collected at the same time the lentil lines were sampled (M14, N = 48). We aligned these data to our new genome assembly and called SNPs as described above, but only considered the 21,342 SNPs already identified from the L14 data set. 18,637 of these SNPs were found to also be variable in the L1–L3 data set.

We used a hierarchical Bayesian model to estimate the allele frequencies for the 21,342 SNPs in L14 (and L11) at each sampled generation, and for the 18,637 SNPs in the L1, L2 and L3 data set (Gompert & Messina, 2016). This model jointly infers genotypes and allele frequencies while accounting for uncertainty in each due to finite sequence coverage and sequence errors, and thereby allows precise and accurate estimates of allele frequencies with low to moderate sequence coverage for individual beetles (see “Allele frequency model” in the OSM for details; Buerkle & Gompert, 2013). Allele frequency estimates were based on two Markov-chain Monte Carlo runs per sample (i.e., line by generation combination), with each consisting of a 5000 iteration burn-in and 15,000 sampling iterations with a thinning interval of 5. We then calculated the mean expected heterozygosity (across SNPs) and pairwise genotypic linkage disequilibrium (measured with *r*^2^ and based on the mean of the posterior for each genotype) among all pairs of SNPs each generation as summary metrics of genetic variation (calculations were made in R without any specialized packages; see DRYAD doi:10.5061/dryad.0tr36fp for the specific code we used).

### Parameterizing and testing a null model of genetic drift

We estimated the variance effective population size (*N_e_*) during the L14 experiment from patterns of allele frequency change, and then used the estimates of *N_e_* to parameterize and test a null model of evolution solely by genetic drift. We did this not as a formal test for selection, but rather to identify the set of SNPs that were most likely to have been affected, at least indirectly (i.e., through linkage disequilibrium), by selection. We estimated variance effective populations sizes as described in Gompert (2016) using a Bayesian bootstrap method (see “Bayesian bootstrap” in the OSM for details; Jorde & Ryman, 2007; Foll *et al*., 2015). Distinct estimates of *N_e_* were obtained for the following generation intervals and (sub)lines: from L14 P to L14 F4, from L14 F4 to L14A F16, and from L14 F4 to L14B F16. We placed a uniform prior on *N_e_* (lower bound = 5, upper bound = 2000), and generated samples from the posterior distribution using 1000 bootstrap replicates. The variance effective population size for the failed L11 line was estimated in a similar manner (see “The L11 line” in the OSM).

We then asked whether the magnitude of allele frequency change for each SNP deviated from null expectations under a model of pure drift, given the estimated values of *N_e_* (we used the posterior median for this, see “Bayesian bootstrap” in the OSM). As with our estimates of *N_e_*, we separately tested for deviations from neutrality for the following generation intervals and (sub)lines: from L14 P to L14 F4, from L14 F4 to L14A F16, and from L14 F4 to L14B F16. We calculated the probability of the observed allele frequency change from the start to end of each of these intervals based on a beta approximation to the basic Wright-Fisher model (Ewens, 2012). Specifically, we assumed *p_t_|p*_0_ ∼ beta(*α* + 0.001, *β* + 0.001), where 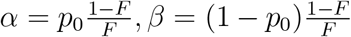, *p*_0_ and *p_t_* are the allele frequencies at the beginning and end of the interval, 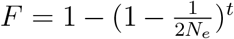, *t* is the number of generations between samples, and *N_e_* is the variance effective population size. We retained SNPs with allele frequency changes more extreme than the 0.1th or 99.9th percentiles of the null distribution for any of the three time intervals for further analyses (Figs. S2, S3). We identified 198 SNPs (188 of which were variable in L1, L2 and L3) based on these relatively conservative criteria, and we hereafter focus primarily on the evolutionary dynamics at and effect of selection on these “focal” SNPs (this is a greater number of SNPs than expected by chance under the null hypothesis of no selection on any SNPs; binomial probability, expected = 128.1 SNPs, *P* = 2.77*e*^−8^).

### Quantifying patterns of linkage disequilibrium over time

To assess the potential for evolutionary independence among these focal loci, we calculated the squared correlation (*r*^2^) between genotypes for all pairs of the 198 SNPs as a metric of linkage disequilibrium (LD). We used Bayesian point estimates of genotypes (posterior means) for this analysis. Estimates of LD were made for each generation and (sub)line and were compared across generations. Hierarchical clustering and network-based methods were then used to identify and visualize groups or clusters of SNPs in high LD, with a focus on patterns of LD in L14–P, L14–F1, L14–F4, L14A–F16 and L14B–F16. We used the Ward agglomeration method implemented in the `R hclust` function for hierarchical clustering (from `fastcluster` version 1.1.24; Müllner *et al*., 2013). Clusters of high LD SNPs were then delineated using the `cutreeDynamic R` function (version 1.63-1) with the cut height set to 99% of the truncated height range of the dendrogram (Langfelder *et al*., 2016). Next, we visualized patterns of LD using networks with each of the 198 SNPs denoted by a node and edges connecting SNPs in high LD. To do this, we created an adjacency matrix from each LD matrix. SNPs were considered adjacent, that is connected in the network, when the *r*^2^ metric of LD was 0.25 or greater; this cut-off corresponds with the 97.5th percentile of the empirical LD distribution for the focal SNPs in L14 P. The `R` package `igraph` (version 1.2.1) was used to construct and visualize these networks (Csardi & Nepusz, 2006).

### Estimating selection in the L14 lines

We estimated the selection experienced by each of the 198 SNPs in L14 from generation P to F4, and then in each subline from generation F4 to F16. These estimates, including their consistency between earlier (up to F4) and later (from F4 to F16) stages of evolutionary rescue (i.e., adaptation to lentil) were used as our primary process-based metric of evolutionary dynamics (patterns of LD and allele frequency changes themselves provided pattern-based metrics of evolutionary dynamics). Comparisons of selection coefficients between L14 sub-lines and time periods allowed us to assess the consistency (over generations and between sublines) of population genomic patterns associated with evolutionary rescue on lentil in *C. maculatus*.

We used approximate Bayesian computation (ABC) to fit Wright-Fisher models with selection and thereby estimate selection coefficients for each SNP in each (sub)line and time period (Ewens, 2012; Gompert & Messina, 2016). This approach uses a stochastic, process-based model of evolution to estimate selection from temporal patterns of allele frequency change while accounting for genetic drift. Thus, in contrast to alternative analytical approaches that do not model drift (e.g., Burke *et al*., 2010), the approach we used does not rely on parallel patterns of change in replicate lines to distinguish between selection and drift. See Fig. 2 for a graphical overview of the model and inference procedure.

**Figure 2:**
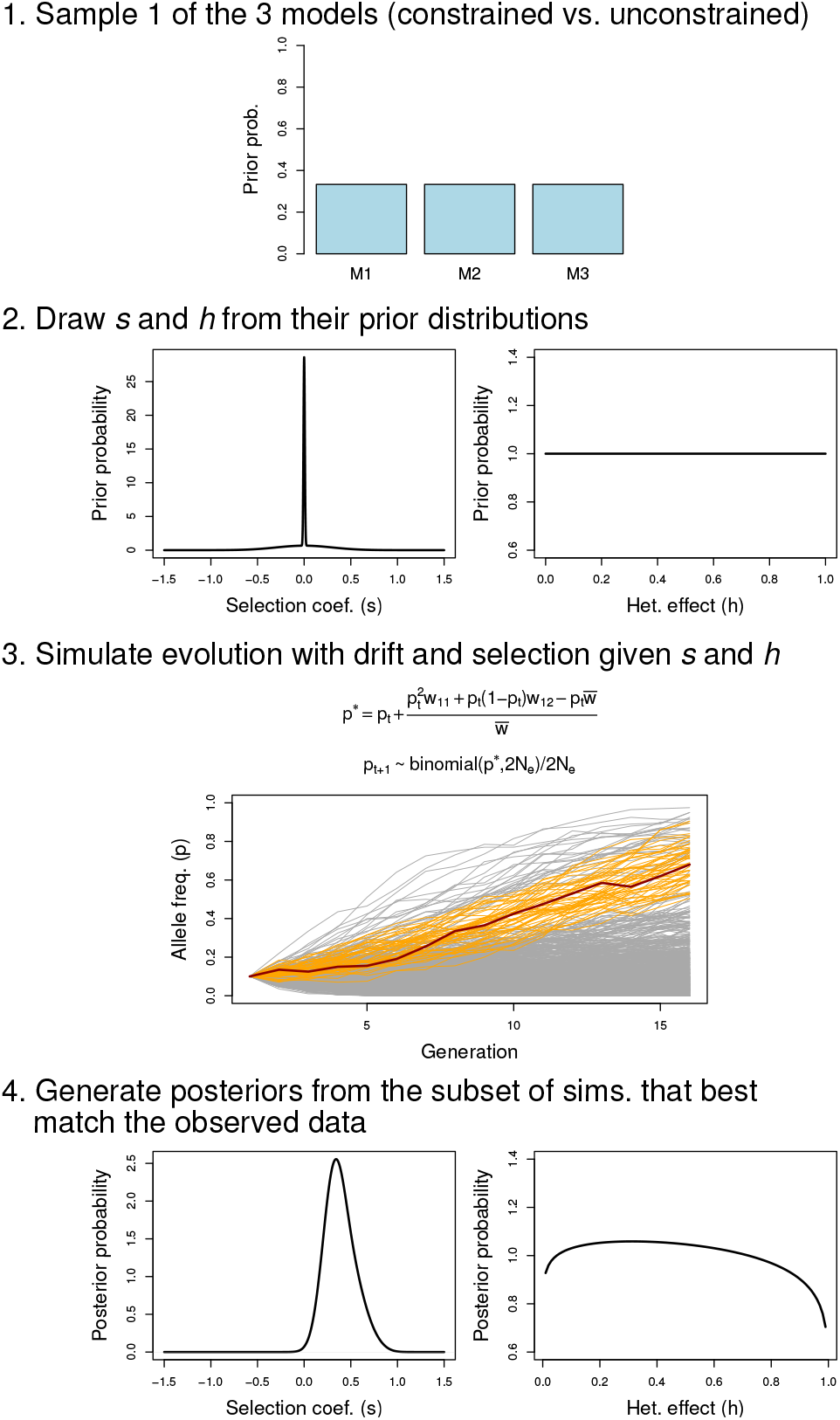
Diagram summarizing the ABC methods used. For each SNP and ABC simulation, we sampled one of the three models of constraint (step 1) and values of *s* (selection) and *h* (heterozygote effect) based on their prior probabilities (step 2). Depending on the model, the same values of *s* and *h* were used across generations or sublines, or different values were sampled for early versus later generations or for different sublines. We repeated this process five million times (per SNP), and each time we simulated evolution under the Wright-Fisher model with selection given the sampled values of *s* and *h* (and our estimates of *N_e_*) (step 3). Here, the first equation gives the expected value of the allele frequency in generation *t* +1 (denoted *p^*^*) based on the current allele frequency (*p_t_*) and fitness values (which are determined by *s* and *h*). The second equation denotes the stochastic component of the Wright-Fisher model, which involves random sampling (drift) around the expected value. The graph below the equations shows 5000 evolutionary trajectories. Orange lines denote the subset that best matched hypothetical observed data for a SNP (shown in red). Values of *s* and *h* from the subset of simulations that best fit (match) the observed allele frequency trajectory are retained and form the basis of the posterior distribution (following correction by local linear regression) (step 4).

We assumed that marginal relative fitness values for the three genotypes at each locus were given by *w*_11_ = 1 + *s*, *w*_12_ = 1 + *hs*, and *w*_22_ = 1, where *s* is the selection coefficient, *h* is the heterozygote effect, and 1 and 2 denote the reference and non-reference allele, respectively. Critically, *s* reflects the combined effects of indirect and (possibly) direct selection on each SNP, and is thus the marginal selection on each SNP. That is, it includes the effect of selection transmitted to a SNP because of LD with one or more causal variants (Gompert *et al*., 2014a; Egan *et al*., 2015; Gompert *et al*., 2017). Our primary interest was in estimating *s*, but we included *h* as a free parameter to account for the effect of uncertainty in *h* on inference of *s*, and to extract any information available from the data on *h*. We considered three evolutionary models with different assumptions about variation in *s* and *h*, (i) a fully constrained model with constant *s* (and *h*) over time and across sublines, (ii) a partially constrained model that allowed *s* and *h* to change at the F4 generation but with identical selection in both sublines, and (iii) an unconstrained model with *a priori* independent values of *s* and *h* before the subline split and in each subline after the split.

With our ABC approach, we first sampled an evolutionary model and values of *s* and *h* from their prior distributions and then simulated evolution forward in time from the parental generation of L14 to generation F16 in sublines A and B while allowing for genetic drift (which was parameterized by the relevant estimate of *N_e_*) and selection (this combines equation 1.24 from Ewens, 2012 with binomial sampling for genetic drift; see details below and in Fig. 2). We assigned a prior probability of 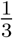 to each model of constraint (i.e., consistency) in selection over time and between sublines (Fig. 2). We assumed a prior distribution on selection (*s*) that was a 50:50 mixture of two Gaussian distributions both with a mean of 0, but one with a modest standard deviation of 0.3 and one with a very small standard deviation of 0.007 (the latter was 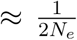 in the post-bottleneck lentil line). This is akin to a spike-and-slab prior, and allows for moderately intense selection while still conservatively putting most of the prior density for *s* near 0 (as in, Gompert & Messina, 2016). The heterozygote effect was assigned a uniform prior over the interval (0,1) (see “Sensitivity to model assumptions” for an assessment of the sensitivity of our results to prior assumptions).

In the ABC simulations, the expected allele frequency (due to selection) in each subsequent generation, *t* + 1, was calculated as 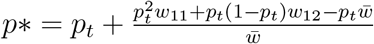, where 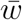 denotes the mean fitness of the population. We then accounted for genetic drift around this expectation by sampling *p_t_*_+1_ ∼ binomial(*p^*^*, 2*N_e_*)*/*2*N_e_*. ABC simulations were implemented in our own computer program (`wfabc-dyn`, version 0.1) written in `C++` using the Gnu Scientific Library (Gough, 2009). Simulation output comprised the full vector of allele frequencies across generations and sublines, which we then compared to the analogous allele frequency vector containing the observed data for each locus. As is standard with ABC methods, posterior distributions for *s* and *h* were generated by retaining (and correcting, see below) the set of parameter values that best recreated the observed allele frequency vector.

We based inferences of *s* and *h* for each of the 198 SNPs on five million simulations. The non-reference allele frequency for each SNP in the L14 founder generation (P) was used to initialize each simulation. We retained the sampled parameter values from the 0.02% of simulations (1000 samples, which provides a reasonable amount of information about the posterior distribution) that generated allele frequency vectors with the smallest Euclidean distance to the observed allele frequency vector (across sublines and generations). We then corrected these sampled parameter values by adjusting them towards the true posterior distribution using a weighted local linear regression (Beaumont *et al*., 2002). This was done with the `abc` function in the `R abc` package (version 2.1) (Csilléry *et al*., 2012). Model posterior probabilities were calculated using a simple rejection method, and posterior probabilities of *s* and *h* integrated over uncertainty in the best model except where noted otherwise. Simulations were used to assess the precision and accuracy of selection coefficient estimates with our ABC framework (see “Evaluation of the ABC approach” and Figs. S4 and S5 in the OSM).

Estimates of *s* were designated as credibly different from zero when the 95% equal-tail probability intervals (ETPIs) of the relevant posterior distribution did not overlap zero. Cases where this was not true do not constitute evidence of neutral evolution, but rather indicate that we cannot confidently distinguish among three possibilities: neutral evolution, selection favoring the non-reference allele, and selection favoring the reference allele. Comparisons of selection coefficients between sublines or time intervals were used to measure the consistency of selection during the L14 evolutionary rescue event and were made by calculating Pearson correlation coefficients (*r*). Rather than basing these calculations on the point estimates of *s*, we obtained posterior distributions for *r* by integrating over uncertainty in *s* (i.e., by calculating *r* for each posterior sample of *s*). Thus, uncertainty in *s* was propagated to downstream summary analyses.

### Estimating genomic change and selection in lines L1, L2, L3 and L11

We quantified patterns of genomic change and selection in the long-established lentil lines (L1-L3) and in the new line that failed to undergo evolutionary rescue and establish on lentil (L11), and compared these to patterns of change and selection in L14. Because of the idiosyncratic nature of these comparisons (L1-L3 were analyzed long after rescue, and L11 failed to adapt), it was not possible to measure the repeatability of genome-wide evolutionary dynamics during rescue. In other words, we could not assess the repeatability of rates or temporal patterns of genomic change, or of the intensity of selection on individual loci. Rather, we made these comparisons to ask whether the outcome of lentil adaptation in L1, L2 and L3, and/or initial patterns of change in the doomed L11 line mirrored the patterns of genomic change and selection (in terms of direction and relative, not absolute, intensity) observed during evolutionary rescue in L14. An affirmative answer would suggest a degree of repeatability for the dynamics of genomic change during evolutionary rescue (or, even more generally, when using lentil regardless of whether rescue is successful), and thus could serve as a basis for future work. Our analyses of L1-L3 and L11 focused on the 198 focal SNPs from the L14 line (only 188 of these were variable in L1-L3, and thus only that subset was considered for those lines).

Selection coefficients were estimated in the long-established lentil lines (L1-L3) using a modified version of the method described above. First, since the mung-bean source line was sampled contemporaneously with the long-established lentil lines rather than at the point in time when the lentil lines were founded, we first simulated evolution by genetic drift backwards in time from the contemporary M14 sample to the mung-bean source population of each lentil line. We used this as a starting point for forward-in-time simulations of evolution by selection and drift in each lentil line (this assumes neutral evolution in the source south Indian line on mung only, not in the lentil lines; see “Alternative ABC models” in the OSM and Gompert & Messina, 2016 for additional details). Variance effective population sizes from Gompert & Messina (2016) were used for these simulations. Values of *s* and *h* were sampled from their prior distributions and the 0.02% of simulations that best matched the observed data were retained as described for L14, but in this case we compared only the final allele frequency in L1 F100, L2 F87 and L3 F85 with the simulated value after 100, 87 or 85 generations of evolution (we lack genetic data from the early stages of adaptation in these lines). Because this constraint greatly reduced the dimensionality of the summary statistics, many simulations gave exact matches to the observed data. This result caused the local linear regression to fail, but also made this correction unnecessary. Hence, we used simple rejection to obtain the posterior distributions of *s* for L1, L2 and L3.

We similarly focused on the final allele frequency for the L11 line, which in this case was in the F4 generation (see “Alternative ABC models” in the OSM for details). However, we were able to use a contemporary sample from the south Indian line (the L14 founders) as the source for L11, and thus did not perform the backwards in time simulation of drift (this sample was taken from the south Indian line within a few months of the time that L11 was founded, and thus any drift would be negligible). Comparisons of selection coefficients across lines (L1-L3, L11, and L14) were then used to assess the repeatability of genomic change on lentil, and were made by calculating Pearson correlation coefficients (*r*) as described in the preceding section.

## Results

### Fitness assays

Survival from egg hatch to adult emergence from lentil seeds was low as expected in the source mung bean population (∼ 1%) (Fig. 1c). Yet survival had risen to 69.2% by the F5 generation. Subsequent to the subline split, survival assays were only conducted in subline A. At generation F10, survival had further increased to 90.2%, and remained high (91.8%) at the F20 generation (subline A). This pattern of rapid adaptation thus closely resembled those observed earlier in the L1-L3 lines (Fig. 1c; Messina *et al*., 2009b).

### Patterns of allele frequency change and LD in the L14 line

We observed substantial evolutionary change over the course of the experiment, with an average net allele frequency change between generations P and F16 of 0.155 in subline A (SD = 0.150) and 0.159 in subline B (SD = 0.155). Average expected heterozygosity also declined over time, from 0.274 (SD = 0.169) in generation P to 0.246 (SD = 0.183) in generation F4, and finally to 0.222 (SD = 0.187; subline A) or 0.220 (SD = 0.174; subline B) in the F16 generation (all standard errors ≈ 0.001) (Fig. S6). Consistent with the observed decline in diversity and the census population bottleneck, the variance effective population size was quite low initially (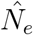 for P to F4 = 8.82, 95% equal-tail probability intervals [ETPIs] = 8.60–9.04; Table 1). Variance effective population sizes then increased between generations F4 and F16 to 68.92 (95% ETPIs = 66.69–71.05) and 56.77 (95% ETPIs = 55.25–58.35) in sublines A and B, respectively. Even in the parental generation, LD was high between nearby SNPs (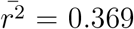 for SNPs <100 bp apart), and modest out to 500 kb 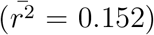 (Table S1, Fig. S7). On average, LD increased over the course of the experiment, although the upper percentiles of the LD distribution reached their maximum by the F4 generation before declining in both sublines.

**Table 1:** Bayesian estimates of variance effective population sizes (*N_e_*) for different sublines and time periods. ETPI = equal-tail probability interval.

| Time period | $N_e$ (median) | $N_e$ (95% ETPIs) |
| --- | --- | --- |
| P-F4 | 8.82 | 8.60–9.04 |
| F4-F16A | 68.84 | 66.69–71.05 |
| F4-F16B | 56.77 | 55.24–58.35 |
| P-F16A | 28.69 | 28.00–29.34 |
| P-F16B | 27.25 | 26.68–27.91 |

Considerably greater evolutionary change was observed for the 198 SNPs with significant deviations from the null genetic drift model (i.e., the focal SNPs). For these SNPs, the average net allele frequency change over the experiment (from P to F16) was 0.611 in subline A (range = 0.004–0.973) and 0.616 in subline B (range = 0.018–0.980) (Figs. 3, S8, Table S2). Many of these SNPs exhibited substantial allele frequency change in a single generation, with a mean (across SNPs), maximum single-generation change of 0.446 (range across SNPs = 0.175–0.745). For 70.7% of these SNPs the maximum change occurred between the F2 and F3 generation (the mean absolute change for this generation was 0.370). By the F16 generation, the initially rarer allele (i.e., the minor allele) had reached a frequency of > 0.90 at 64.1% of these SNPS, and > 0.98 for 29.2% (subline A) or 22.2% (subline B) of them. Frequency changes during the first four generations were only modestly correlated with changes after the formation of the the two sublines (*r_P−F_* _4*,F* 4*−F* 16*A*_ = 0.127, 95% confidence interval [CI] = −0.012–0.262; *r_P−F4,F4−F16B_* = 0.234, 95% CI = 0.098–0.362), while evolutionary changes between sub-lines after the split showed high levels of parallelism (*r_F_* _4_*_−F_* _16*A,F* 4*−F* 16*B*_ = 0.743, 95% CI = 0.674–0.800; Fig. S9).

**Figure 3:**
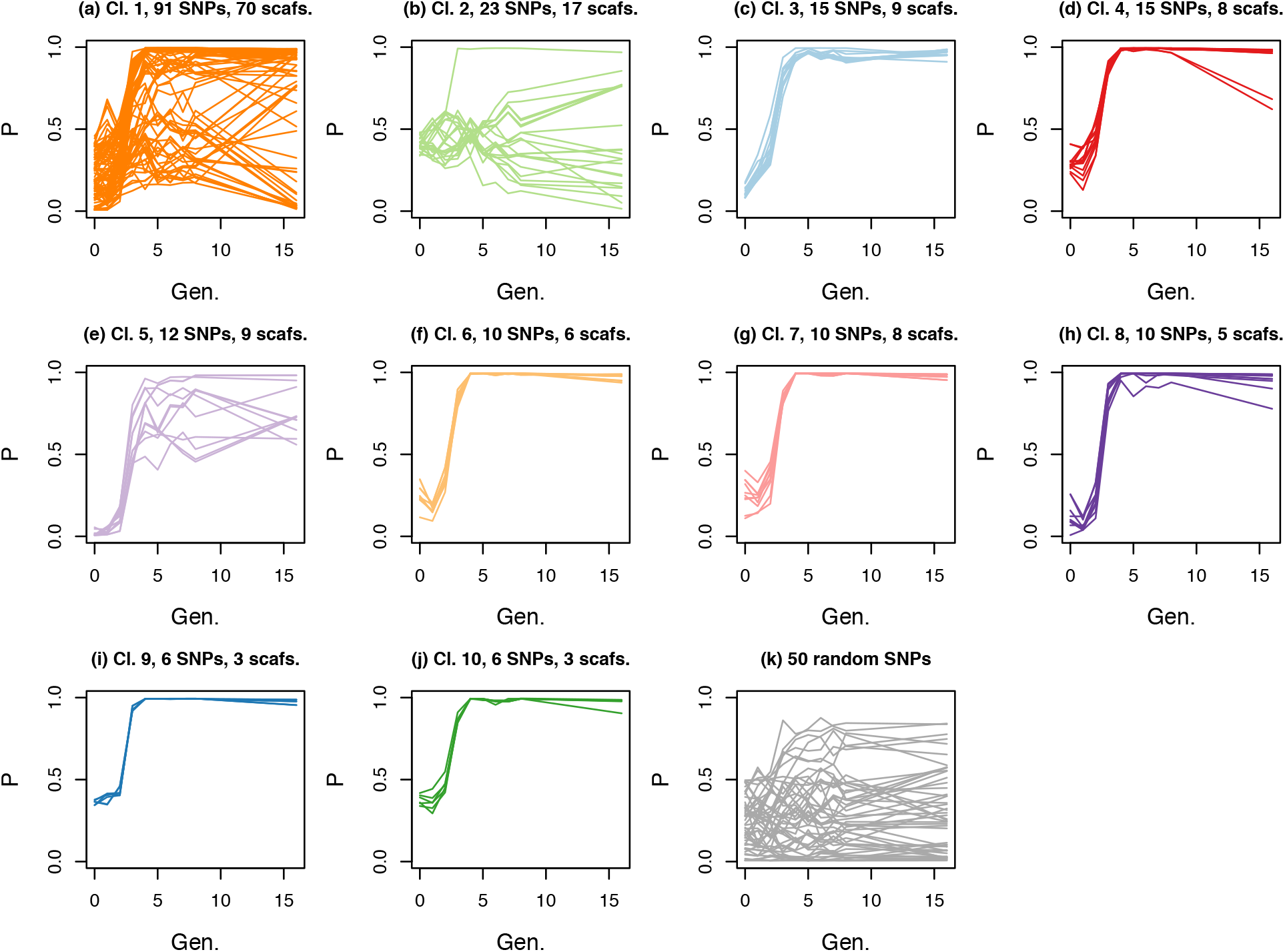
Patterns of allele frequency change for L14 subline A (L14A). Panels (a)–(j) show allele frequency (*P*) over time (Gen. = generation) for the 198 focal SNPs. Each line shows the allele frequency trajectory for a single SNP and these are organized into panels by the LD clusters delineated in the F1 generation (Cl. = cluster number; see Fig. 4 and the main text for details). Colors correspond with those from L14–F1 in Fig. 4(a). The number of SNPs and number in each panel and number of scaffolds on which they reside is given. Panel (k) shows patterns of change for 50 randomly selected SNPs. In all cases, the frequency of the minor allele from the parental generation is shown. See Fig. S8 for similar results from L14B.

The 198 focal SNPs did not evolve independently, but instead were organized into clusters of high LD loci that exhibited similar patterns of allele frequency change (Figs. 3, 4, S10). We identified 16 and 10 clusters of high LD SNPs in the L14–P and L14–F1, respectively, which were reorganized into six high LD clusters by the F4 generation. LD within clusters was considerably higher than LD among clusters (e.g., mean *r*^2^ within, 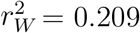, versus mean among, 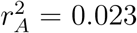 in L14–F4; Fig. 4). Despite the fragmented nature of our reference genome (Fig. S1), we found that cluster membership was consistent with physical proximity, such that SNPs on the same scaffold were more likely to be assigned to the same cluster (*p <* 0.001 based on a randomization test in L14–F1). With that said, patterns of LD and cluster membership shifted over the experiment, particularly during the first four generations (Fig. 4b), such that pairwise LD in generations F1 and F4 were only modestly correlated (*r_F_* _1*,F* 4_ = 0.199). Patterns of LD changed less after that; the correlations in pairwise LD between F4 and L14A–F16 and L14B–F16 were *r_F4,F16A_* = 0.605 and *r_F4,F16B_* = 0.569, respectively.

**Figure 4:**
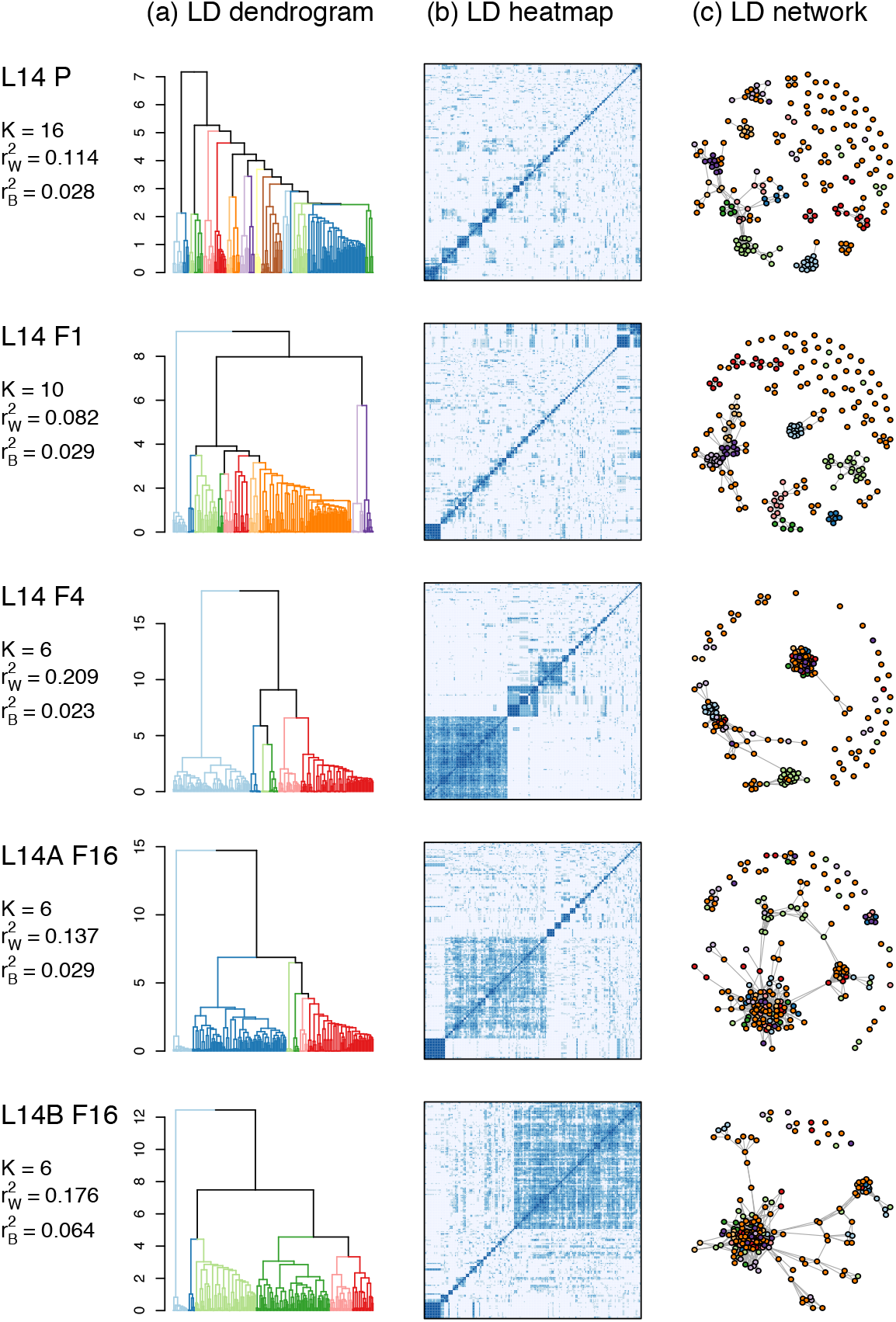
Patterns of LD among the 198 focal SNPs for L14–P, L14–F1, L14–F4, L14A–F16 and L14B–F16. Panel (a) shows dendrograms from hierarchical clustering of SNPs based on LD, with colors denoting clusters delineated with the `cutreeDynamic` function (colors do not track clusters across generations). The number of clusters (*K*) and mean LD for SNPs in the same 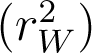 versus different 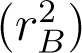 clusters are given. The corresponding pairwise LD matrixes are shown as heat maps in panel (b) (darker shades of blue denote high LD). Panel (c) shows networks connecting SNPs (nodes = colored dots) with high LD (r^2^ ≥ 0.25). Nodes are colored based on their cluster membership as defined by hierarchical clustering in the F1 generation (see panel a) (compare to Fig. S10 where, for each generation, nodes are colored based on their cluster membership in that generation).

### Strength and consistency of natural selection in the L14 line

For most SNPs, constrained and unconstrained models of selection had similar posterior probabilities (Fig. S11). Consequently, rather than focus on a specific model, we report model-averaged selection coefficients. Consistent with the observed patterns of allele frequency change, selection coefficients were large on average, especially during the early stages of adaptation (i.e., from L14–P to L14–F4) (allele frequency change and estimates of selection were strongly correlated, with *r>* 0.8; Fig. S12). In particular, the average intensity of selection was 0.388 in L14 from P to F4, and 0.207 and 0.211 in sublines A and B between the F4 and F16 generations (Fig. 5) (see Figs. S13, S14, S15, S16, S17 and S18 and text in the OSM for results using different priors). Of these 198 SNPs, we detected a credible effect of selection (that is, 95% ETPIs for *s* not overlapping zero) in 53 SNPs from six of ten LD clusters during the early phase of adaptation (from P to F4), and 53 and 51 SNPs from four of ten LD clusters during the later stage of adaptation (F4-F16) in sublines A and B, respectively (here we define LD clusters based on patterns of LD in L14–F1; some but not all of these clusters had a credible effect of selection in both early and later stages of adaptation; Fig. 6). Nearly all credible estimates of *s* were negative, implying selection for the non-reference allele, which was generally the minor allele (Fig. 5). Estimates of *h* (the heterozygote effect) were associated with considerable uncertainty, but there was a slight signal of an overall negative correlation between *s* and *h* (see “Heterozygous effect”, Table S3 and Figs. S19 and S20 in the OSM for details).

**Figure 5:**
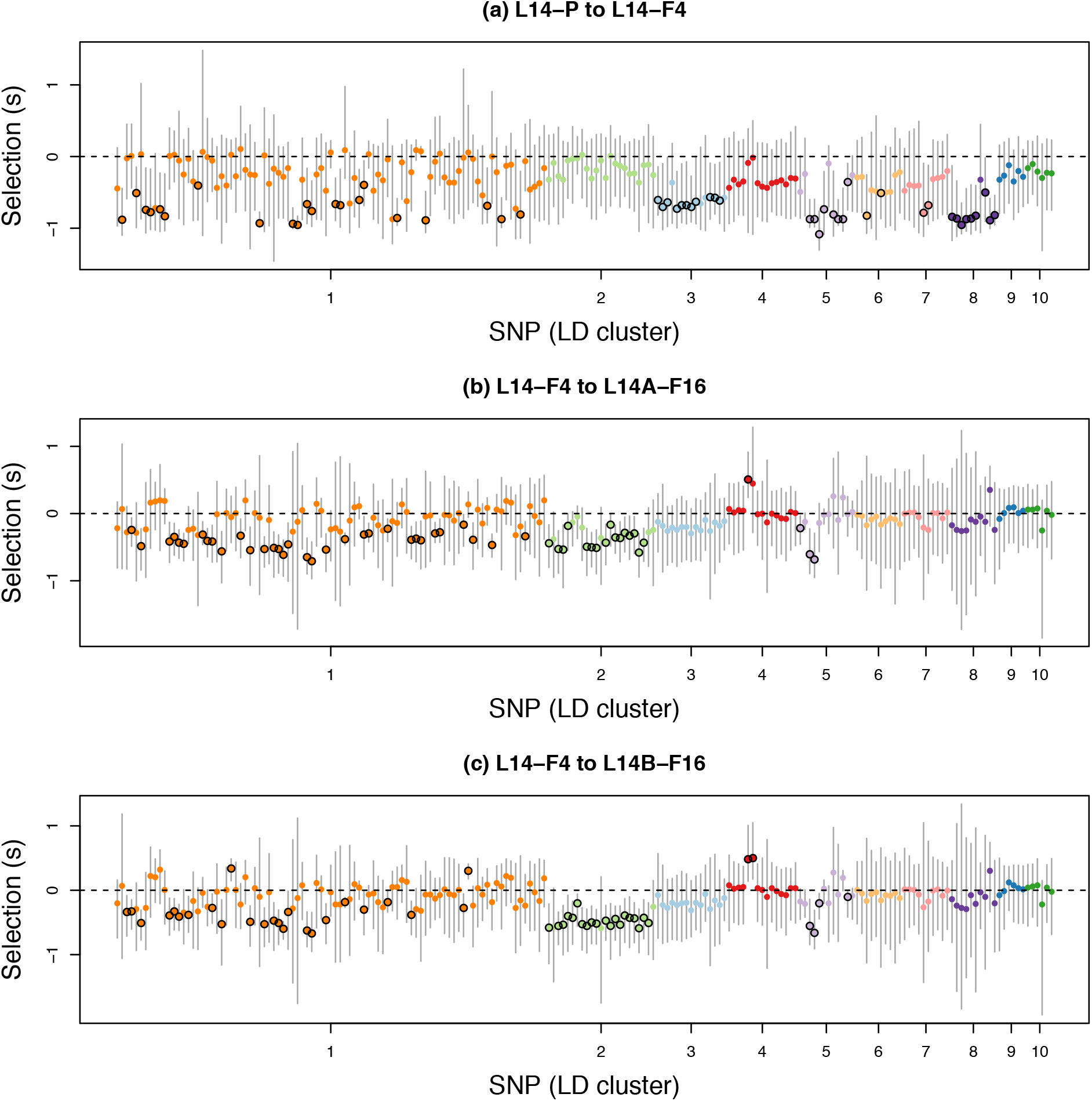
Plots show Bayesian estimates of selection coefficients for the 198 focal SNPs for (a) the origin of L14 through the F4 generation (L14–P to L14–F4), (b) the F4 generation to the F16 generation in subline A (L14–F4 to L14A–F16), and (c) the F4 generation to the F16 generation in subline B (L14–F4 to L14B–F16). Dots and vertical bars denote posterior medians and 95% equal-tail probability intervals (ETPIs), respectively. Colors and the order of SNPs reflect LD cluster membership in the F1 generation. Black circles around dots denote cases where the 95% ETPIs exclude 0. For the purpose of visualization, we have polarized estimates of *s* such that negative values indicate selection favoring the minor allele.

**Figure 6:**
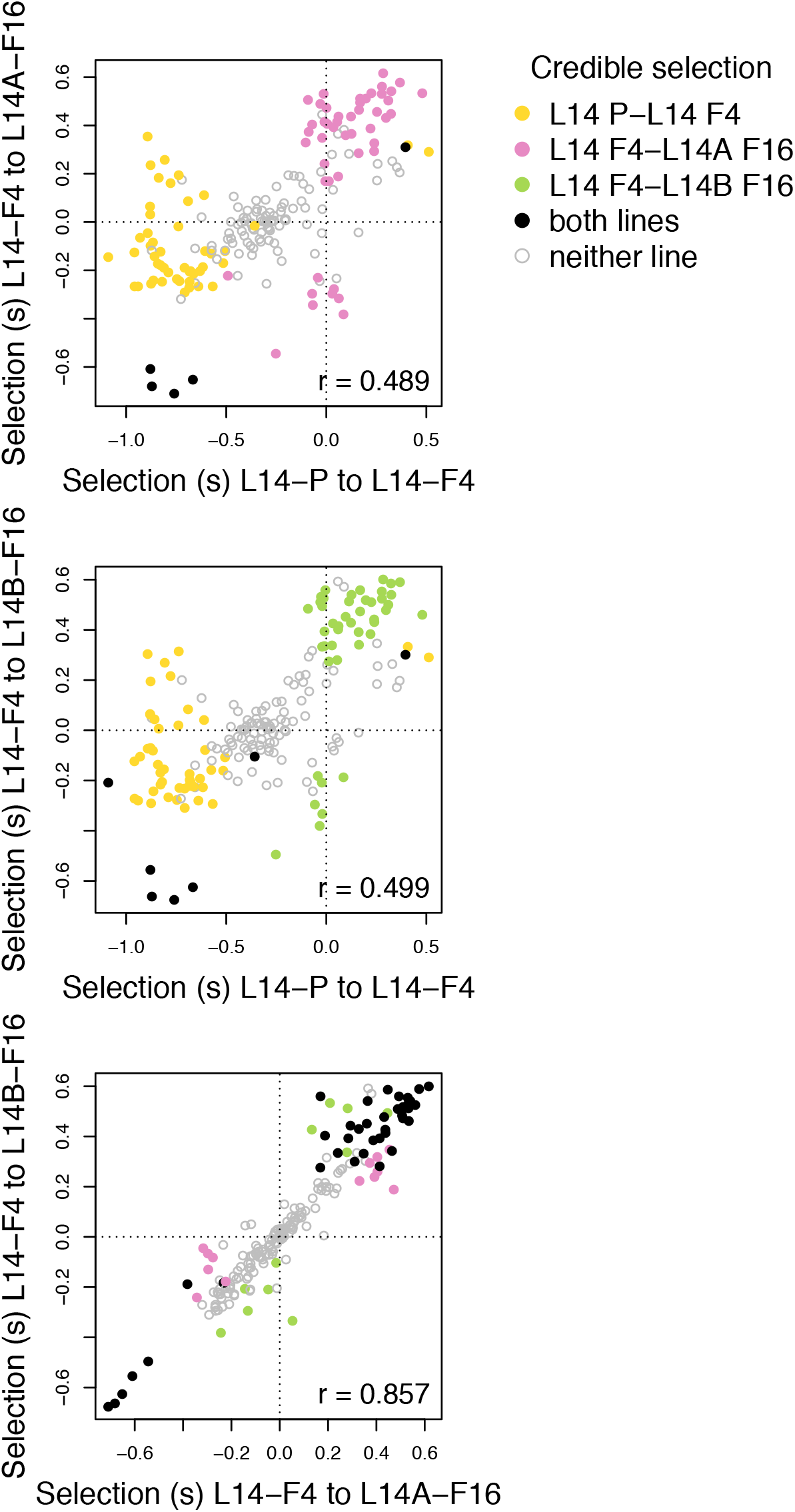
Scatter plots show associations between selection coefficient estimates for the 198 focal SNPs in L14 for different time intervals and sublines. Dots correspond to SNPs and are colored based on whether there was credible evidence of selection in each subline/interval. Pearson correlations account for uncertainty in estimates of selection (i.e., they are not based solely on the point estimates shown here).

Only five and seven SNPs had credible effects of selection during both time periods for sublines A and B, respectively (Fig. 6a,b). Nevertheless, estimates of *s* during early (between P and F4) and late (from F4 to F16) adaptation were moderately correlated (*r_P−F_* _4*,F* 4*−F* 16*A*_ = 0.489, 95% ETPIs = 0.373–0.587; *r_P−F4,F4−F16B_* = 0.499, 95% ETPIs = 0.387– −0.592) (Table S4). Moreover, we never detected credible effects of selection with opposite signs between time periods. We obtained similar results when we based our inferences only on the fully unconstrained model (see “Sensitivity to model assumptions” and Figs. S18 and S21 and Table S5 in the OSM for details). We detected much greater consistency in estimates of *s* between the two sublines during the later stages of adaptation (*r_F_* _4_*_−F_* _16*A,F* 4*−F* 16*B*_=0.857, 95% ETPIs = 0.753–0.914; Fig. 6c) than between time periods. Forty SNPs had credible effects of *s* in both sublines, and always with the same sign.

### Comparisons with other lines

On average, estimates of *s* were lower for the long-established lentil lines with means of 0.067, 0.103 and 0.022 in L1, L2 and L3, respectively. Lower estimates of *s* are expected, as patterns of change were averaged over longer periods of time and thus weaker selection over a longer period of time could explain the observed changes (this effect was evident in Gompert & Messina, 2016). Similar numbers of SNPs had values of *s* credibly different from zero (43 in L1, 55 in L2, and 10 in L3). Correlations in selection coefficients among the three long-established lines ranged from 0.094 to 0.262, and were thus considerably lower than correlations in selection between L14 sublines or time intervals (Fig. 7, S22). There was an even weaker association between selection in the L14 and any of the long-established lentil lines, with correlations in selection ranging from −0.024 to 0.050 (Table S4). Correlations in patterns of allele frequency change among L1, L2 and L3 and between these lines and L14 were generally consistent with correlations in selection coefficients, with moderate correlations in change among L1-L3 (*r*=0.366– −0.653), and much weaker correlations between L14 and these lines (*r* =-0.071– −0.123) (Fig. S9).

**Figure 7:**
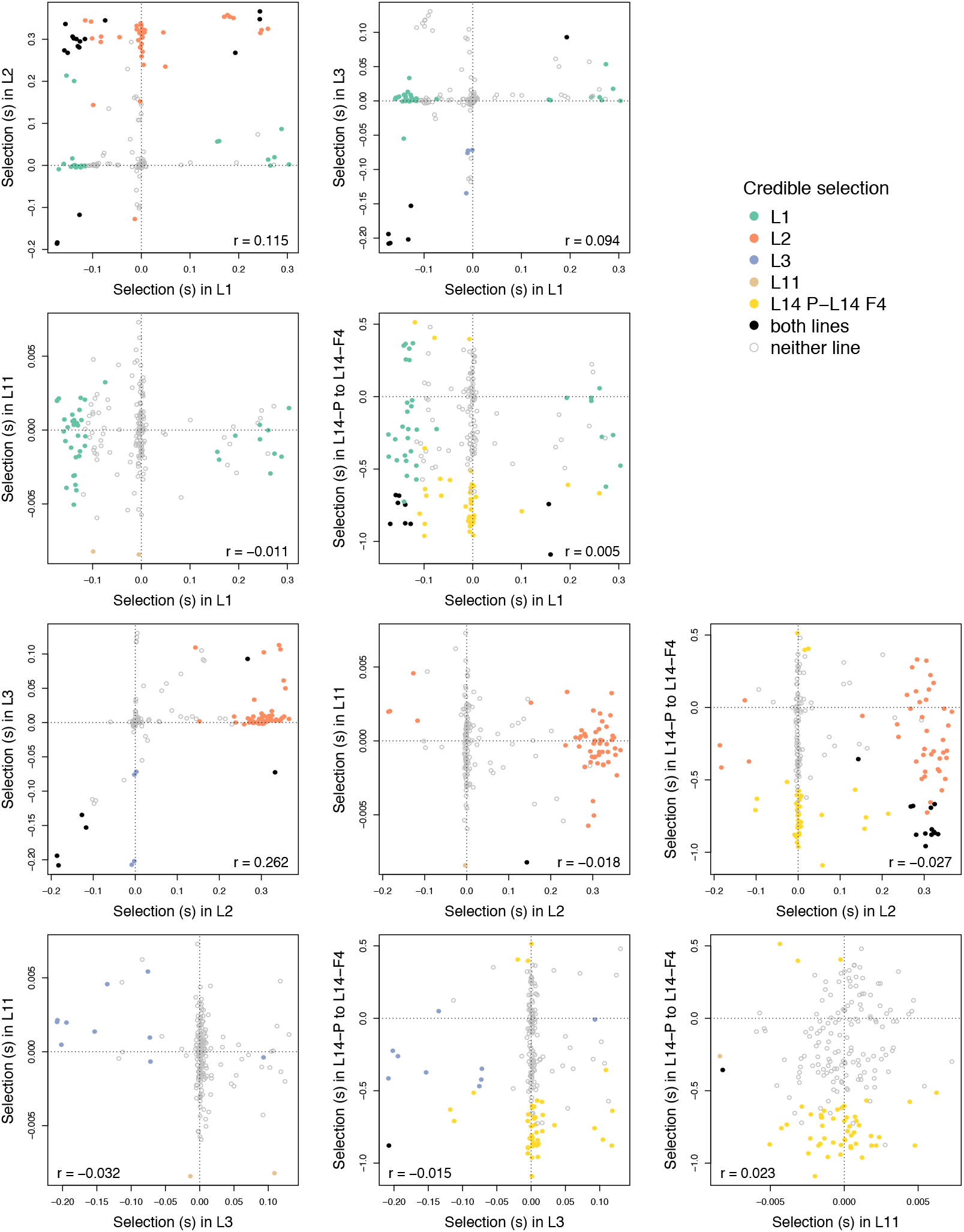
Scatter plots show associations between selection coefficient estimates for the focal SNPs across lines. Results are shown here for all comparisons involving the early stage of rescue in L14 versus other lines. For comparisons with lines L1, L2, and L3 the 188 SNPs present in those lines are shown (a single point for L11 was omitted for visualization as it had an extreme but not-credible estimate of *s*). Dots correspond to SNPs and are colored based on whether there was credible evidence of selection in each (sub)line. Pearson correlations account for uncertainty in estimates of selection (i.e., they are not based solely on the point estimates shown here).

Compared to the L14 line, we detected much less evolutionary change genome-wide in L11 from generation P to F4, with a mean absolute change of 0.053 (SD = 0.050; mean heterozygosity in L11 F4 was 0.276). Consistent with the lack of genomic change, the variance effective population size of this line was considerably higher than it was for L14 during the same time period, *N_e_* = 134.2 (95% ETPIs = 127.2–141.9) (compare to Table 1). This pattern of limited allele frequency change held for the 198 focal SNPs as well (mean change = 0.085, SD = 0.0676), and patterns of allele frequency change at these SNPs were mostly unrelated to patterns of changes in other lines (most *r* = −0.121– −0.093) (Fig. S9). The only exception was for the comparison with change between the L14 founder (P) and L14 F4, with *r_L_*_11_*_,L_*_14_*_P-F4_* = 0.305 (95% CIs = 0.173–0.426). Consistent with the the lack of evolutionary change, we found little evidence of selection affecting the focal SNPs with a mean estimate of *s* = 0.002, and only two SNPs with credible evidence of selection. Likewise, estimates of *s* in L11 were mostly independent of estimates of *s* in the other lines (*r* = −0.032–0.023, with all 95% CIs spanning 0) (Figs. 7, S22, Table S4).

## Discussion

Many important questions in molecular ecology, such as those concerning the evolution of range limits, population persistence in human-altered environment, and the evolution of antibiotic/herbicide/insecticide resistance, focus on cases where adaptation might or might not prevent local extirpation or extinction, that is, cases of potential evolutionary rescue. Thus, a better understanding of not only the factors affecting the probability of rescue, but also of the genetic basis and dynamics (i.e., patterns and rates of change) of evolutionary rescue is broadly relevant to this field. Here, using an evolve-and-resequence approach with fine-scale temporal resolution, we have shown that evolutionary rescue in *C. maculatus* on lentil can occur via rapid evolutionary changes at multiple loci. We observed exceptionally rapid and pronounced evolution during the first four generations on lentil, with an average maximum rate of allele frequency change of 0.446 in a single generation for a set of 198 focal SNPs.

We found evidence of very strong selection on the 198 focal loci during this experiment, with an average intensity of (direct plus indirect) selection on these loci ranging from *|s|* = 0.207 to *|s|* = 0.388 (depending on the subline and time interval) (Fig. 5). Although this magnitude of selection is much stronger than is commonly assumed in population-genetic theory, it is consistent with strong selection detected in other systems, such as sticklebacks (Barrett *et al*., 2008), phlox (Hopkins & Rausher, 2012), flies (Cardoso-Moreira *et al*., 2016) and stick insects (Gompert *et al*., 2014a; Nosil *et al*., 2018), as well as with the observed rapid rise in survival of *C. maculatus* on lentil (Fig. 3). Thus, our work further highlights the importance of developing population-genetic theory with a greater emphasis on strong selection and rapid adaptation, especially in populations that colonize stressful or novel environments (e.g., Gompert, 2016; Messer *et al*., 2016). We also found that different sets of loci were associated with the very early (i.e., F1-F4) versus later (i.e., F4-F16) stages of adaptation to lentil. In other words, patterns of selection varied considerably even over the relatively short time scale of this experiment. In contrast, we observed extreme parallelism (*r* = 0.857) in patterns of selection and evolutionary change in sublines that were separated after the L14 population recovered from an initial bottleneck (Fig. 6c). In the following sections, we discuss these findings in more detail, contrast our core results from L14 with patterns changes in other independently-derived lines (i.e., L1, L2, L3 and L11), and highlight important caveats related to our findings.

### The genetic architecture and evolutionary dynamics of rescue

Survival rates on lentil increased from ∼1% to over 90% in just 10 generations (i.e., survival rates increased ∼90-fold over a short period of time; Fig. 1c). During this time, the new lentil line (L14) went through a severe bottleneck with the variance effective population size (*N_e_*) dropping to fewer than 10 individuals before rebounding. This demographic rebound was likely driven by adaptive evolutionary changes at (i) several to tens of causal loci, or 1. (ii) a similar number of sets of tightly linked mutations with smaller individual effects on fitness (e.g., Linnen *et al*., 2013). In particular, we detected very strong selection (combined direct and indirect) affecting >100 SNP markers, which were organized into 6–16 high LD clusters. We hypothesize that each cluster comprises SNPs in LD with one or more distinct causal variants. If we are correct, our results suggest that rapid adaptation to lentil is driven by strong selection on oligogenic to moderately polygenic variation (as in Orr, 2005 and Bell & Gonzalez, 2009), similar to adaptation to freshwater in marine sticklebacks (Jones *et al*., 2012; Lescak *et al*., 2015). This is consistent with theory predicting a greater role for mutations of large effect (and fewer total genes/gene regulatory regions) during the early stages of adaptation, particularly when a population is far from a phenotypic optimum, as might commonly occur in cases of evolutionary rescue (Orr, 2005; Bell, 2017). However, we will have underestimated the number of causal loci if (i) some causal loci were not in LD with any of our SNPs, or (ii) if multiple causal loci were in high LD with the same SNPs. Consequently, we cannot exclude a more highly polygenic basis for evolutionary rescue. Likewise, because we were conservative in our tests for selection, we likely missed some or even many loci with smaller effects on fitness.

The minor allele at more than 100 SNPs reached a frequency >90% within 16 generations (and in some cases within five generations) (Fig. 3). While we lack data on the underlying causal variants, we can assume that such variants evolved at least this rapidly during the same time period, as direct selection on a causal variant should generally exceed indirect selection on a marker locus in LD with that variant (e.g., Gompert, 2016; Gompert *et al*., 2017). We think this constitutes evidence that selection on standing genetic variation fixed or nearly fixed alleles (or haplotypes) at many of these causal loci. Our results differ from other recent evolve-and-resequence experiments in eukaryotes (mostly involving *Drosophila*) where adaptation occurred by more subtle shifts in allele frequencies and incomplete selective sweeps (Burke *et al*., 2010; Orozco-terWengel *et al*., 2012; Burke *et al*., 2014; Tobler *et al*., 2014; Graves Jr *et al*., 2017) (but see, Michalak *et al*., 2018). For example, in an experiment where flies evolved under novel laboratory conditions with elevated and fluctuating temperatures, SNPs that showed the most pronounced evolutionary change during the first 15 generations of evolution (median change = 0.28), exhibited little allele frequency change after that (median change = 0.03) with only only 9% reaching a frequency >0.9 after 37 generations (Orozco-terWengel *et al*., 2012). Similarly, a highly replicated study of *Drosophila* populations selected for development time found little or no evidence of hard/complete selective sweeps even after >800 generations of evolution (Graves Jr *et al*., 2017).

We think these differences can be explained by the harshness of the experimental environment, and by the related demographic consequences of the imposed environmental shift. When the Indian *C. maculatus* population is shifted onto lentil, absolute mean fitness is initially extremely low, and population decline occurs. In contrast, the experimental environments in the *Drosophila* studies described above (e.g., accelerated culture conditions, altered rearing temperatures, etc.) did not depress absolute mean fitness enough to cause population decline, and thus were arguably more benign. We hypothesize that *C. maculatus* beetles transferred to lentil are far from a fitness peak (relative to the situation in the *Drosophila* studies), such that selection continues to favor the same alleles until they fix (i.e., selection does not transition from directional to stabilizing before fixation). Genetic drift during the initial population bottleneck might have also contributed to the fixation of advantageous alleles during our experiment once they became relatively common.

Despite the constant host environment during the experiment, selection on individual loci varied across generations, particularly in terms of the magnitude (but not direction) of selection. Several complementary explanations may account for this observation. First, given the observed patterns of allele frequency change at the SNP markers, some causal variants likely fixed or nearly fixed within the first five generations. After this, selection on these variants would have ceased, thereby reducing or eliminating selection on linked SNP markers. Second, epistatic interactions could have altered the marginal fitness effects of causal variants as allele frequencies changed. Epistatic interactions have previously been shown to play an important role in adaptation in several species, including mice (Steiner *et al*., 2007), yeast (Ono *et al*., 2017), and bacteria (Arnold *et al*., 2018), and future work mapping the genetic basis of host-specific performance or fecundity traits in adapted/maladapted *C. maculatus* could test for epistasis in this system. Third, direct selection on causal variants could be constant, but indirect selection on our SNP markers could shift as allele frequencies and LD evolve. Given the major shifts we see in patterns of LD, this is almost certainly part of the reason for the variable strength of selection over time. Lastly, some sources of selection could be density dependent. Male-male competition is common in high-density populations of *C. maculatus* (Hotzy & Arnqvist, 2009), and larvae from our Indian source population exhibit particularly strong contest competition within seeds (Messina, 1991; Fox & Messina, 2018).

### Low repeatability of genomic change on lentil

Because we documented the full course of evolutionary rescue in the L14 line (even if adaptation was ongoing, the population had been rescued from extinction), patterns of genomic change in the L14 experiment provide direct insights into the population genetic/genomic processes associated with rescue. In contrast, even non-neutral genetic differences between each of the long-established lentil lines (L1-L3) and the source south Indian population include a mixture of adaptive changes that occurred during rescue and adaptive changes that occurred after rescue was complete, and any adaptive evolutionary change that occurred in L11 was insufficient to rescue the line from extinction. Thus, we cannot ask whether the SNPs that changed most (and thus were most likely affected by selection) in L14 during rescue followed similar dynamics in other lines (e.g., similar rates and directions of change and in the same temporal sequence). However, we were able to ask whether SNPs associated with rescue in L14 showed overall patterns of change in the other lines that were at least consistent with being associated with (partial) adaptation to lentil. An affirmative answer would imply that evolution on lentil is repeatable at the genetic level (at least to a degree and without a focus on dynamics per se), perhaps even in cases where extinction occurs.

This was not what we observed. Instead, total allele frequency change and estimates of selection for the 198 focal SNPs in L1, L2, L3 and L11 were mostly independent of patterns of change and selection in L14 (Figs. 3, 7). The only notable exception was for allele frequency change in L11, which exhibited a modest (*r* = 0.305) but significant correlation with change during the during the first four generations of L14 on lentil (Fig. 3). Thus, there is some, albeit limited, evidence that the evolutionary path L11 followed was not wholly independent of the path followed by L14, even though changes were much larger in the latter and resulted in rescue (we discuss this more below). We also detected greater consistency in patterns of genomic change among the three long-established lentil lines (consistent with Gompert & Messina, 2016), than between any of these and L14. This can perhaps be explained by evolutionary changes within the south Indian mung bean line. Evolutionary changes within the source mung bean line could have likely altered the standing genetic variation initially available for adaptation to lentil in each line. Given the moderately high variance effective population size in this source line (*N_e_* = 1149; Gompert & Messina, 2016) and the fact that the population has been kept on the same host for >1000 generations, we expected minimal evolution within this line. Nonetheless, it is clearly still evolving. L2 and L3 were formed within just a few generations of each other, and L1 was started about 20 generations before that (Messina *et al*., 2009b; Gompert & Messina, 2016).

Interestingly, rates of evolutionary change were much lower in the failed L11 line than over the comparable number of generations in the successful L14 line. Likewise, and despite the fact the census sizes of both lines were both reduced to a few hundred adult beetles, the variance effective population size in L11 (∼134) was considerably higher than in L14 (∼9). This difference suggests that variation in fitness was greater in L14 than in L11 (leading to a lower variance effective population size in L14), and is consistent with the observed pattern of rapid adaptation to lentil in L14 and the lack (or limited nature) of adaptation to lentil in L11. Determining whether such differences in variance effective population size generally distinguish cases where evolutionary rescue does or does not occur will require future work with replicated cases of successful and failed rescue.

## Conclusions

We documented rapid adaptation to a stressful host by seed beetles, and showed that it was associated with exceptionally rapid evolutionary change at numerous loci. This result does not mean that all (or any) of the focal SNPs drove adaptation to lentil. Instead, these SNPs were in LD clusters associated with the actual causal variants and thus were likely affected by selection indirectly. Nevertheless, this study shows how a population can rapidly adapt from standing variation at multiple loci across the genome, resulting in a 45 to 90-fold increase in survival and evolutionary rescue from extinction in as few as 10 generations. By showing what is possible, our results might help explain colonization of extreme or stressful environments in nature (Kawecki, 2008), including rare shifts onto marginal host plants in phytophagous insects such as the occasional use of lentil by *C. maculatus* (Credland, 1987, 1990).

In terms of the evolution of increased survival rates, successful cases of evolutionary rescue of *C. maculatus* on lentil exhibit highly repeatable dynamics (Fig. 1c). However, this was not generally true at the genetic level, with the notable exception of parallel (repeatable) changes in the two L14 sublines (Fig. 6). Similar differences in repeatability at genetic and phenotypic levels has been documented in a variety of systems (Blount *et al*., 2018). Beyond this, some of this variation in repeatability of adaptation to lentil at the genetic level likely stems from differences in shared genetic variation available for selection across these cases (as has also been seen in evolve-and-resequence studies in *Drosophila*; Seabra *et al*., 2017). Because lentil is a very stressful host, each *C. maculatus* line went through a severe bottleneck when it was shifted onto lentil (Gompert & Messina, 2016). Thus, the subset of adaptive genetic variation (or adaptive gene combinations) available for selection in each line was likely quite different (e.g., Charlesworth, 2009; Tinghitella *et al*., 2011), which can limit repeatability at the genetic level. These results suggest that the early stages of adaptation to lentil exhibit chaotic dynamics, in the sense that evolutionary trajectories of alleles during this time period are sensitive to initial conditions (i.e., to the specific adaptive alleles present, their frequencies and patterns of LD) (e.g., Rego-Costa *et al*., 2018), and that stochastic variation in initial conditions (due to the demographic bottleneck) expose this sensitivity. The evolutionary consequences of this sensitivity to initial conditions could be amplified by epistatic interactions among alleles that contribute to early and later stages of adaptation (e.g., Lagator *et al*., 2014). Whether this occurs in *C. maculatus* is unknown, but genetic crosses could be used to detect lentil performance/preference QTL with epistatic effects and thereby test this hypothesis. Finally, even in the absence of bottlenecks, repeatability at the genetic level could be low if many redundant loci offer alternative routes for adaptation to lentil, and this could further explain our results.

In conclusion, our results suggest that demographic history can be a key determinant of the extent of parallel evolution at the genetic level, and that bottlenecks could decrease the repeatability of genomic change in cases of evolutionary rescue by exposing chaos. Consequently, understanding the repeatability/predictability of evolution might require considering both ecological (e.g., demographic) and evolutionary processes and a better integration of eco-evolutionary thinking throughout evolutionary biology (e.g., Hendry, 2016).

## Acknowledgments

This manuscript was improved by comments on earlier drafts by J. Fordyce, L. Lucas, C. Nice, P. Nosil, T. Saley, A. Springer, and several anonymous reviewers. We thank C. Bourgeois, S. Thelen, and C. Willden for technical assistance. This research was supported by the Utah Agricultural Experiment Station, Utah State University (UAES paper number 9119). The support and resources from the Center for High Performance Computing at the University of Utah are gratefully acknowledged.

## Author Contributions

ZG and FJM conceived and designed the study. FJM ran the selection experiment. ZG generated the DNA sequence data. AR and ZG analyzed the data. ZG, AR and FJM wrote and revised the manuscript.

## Data Accessibility

DNA sequence reads are available from NCBI’s SRA (BioProject number PRJNA480050). Other data and computer code used to analyze these data are available from DRYAD (doi:10.5061/dryad.0tr36fp).

