## Supplemental materials for "Dynamics of genomic change during evolutionary rescue in the seed beetle *Callosobruchus maculatus*"

### Supplemental Methods and Results

#### Culturing and establishing the lines

Both the long-established lentil lines and the new line produced for the current study were derived from the same base population of *C. maculatus* that was originally collected from southern India (Messina, 1991; Mitchell, 1991). This population had been continuously reared on mung bean, *Vigna radiata* (L.) (Wilczek), for >300 generations when we formed the new lentil-adapted line. Each generation has been started by adding 1500-2500 newly emerged adults (estimated by volume) to a 2-liter jar containing 750 g of mung beans, or about 12,000 seeds. We reared beetle lines and conducted the phenotypic assays described below in a growth chamber at 24°C and constant light. Three original lentil-adapted lines (L1-L3) were established as described by Messina *et al.* (2009b). Lines were formed by adding >2000 (L1) or >4000 (L2-L3) newly emerged adults to two standard culture jars containing 750 g of lentil seeds. Multiple, previous assays demonstrated that, for the Indian beetle population, initial survival to adult emergence is only 1-2% in lentil, development time is very long (averaging about 60 days), and the body size of the few emerging adults is unusually small. Consequently, there is a severe, initial bottleneck, and more than half of the attempts to produce a self-sustaining, lentil-adapted population eventually fail (Messina *et al.*, 2009b; Gompert & Messina, 2016). In the lines later designated as L1-L3, survival increased rapidly, and we were able to implement standard culturing techniques, with >500 beetles per generation, by the fifth generation. Survival in these lines reached >60% after only five generations, and >80% in fewer than 20 generations (Messina *et al.*, 2009b). At the same time, there were substantial decreases in development time and increases in body size.

We followed the same protocol to establish a new lentil line (L14) so that it would be possible to sample the line for population genomic analyses every generation during the early phase of adaptation. The L14 line was started with > 4000 beetles (split between two jars) from the M1 subline on mung bean. F1 beetles emerged from lentil seeds after 45-70 days, with most emerging after 55-65 days (initial egg-to-adult development time was thus approximately twice as long as it is in a lentil-adapted line). One to two hundred F1 beetles were pooled from the two jars and were transferred to a new jar over several days to form the F2 generation. The number of adults transferred to clean seeds increased gradually, so that the F2–F4 generations were started with at least a few hundred individuals. After five generations, the L14 population size was sufficiently high to implement standard culturing techniques, which involve transferring >2000 beetles to a new batch of 750 g lentil seeds in each subsequent generation.

#### The L11 line

In addition to the four lines described in the previous section, ten more attempts were made between October 2013 and September 2014 to adapt the south Indian *C. maculatus* population to lentil (lines L4–L13). In all ten cases, evolutionary rescue was not observed and the populations failed to establish (they either dwindled toward extinction or never reached sufficient numbers to begin standard culture procedures). Here, we focus on one of these lines, L11, which we sequenced as part of the current project. We formed the L11 line on May 23rd, 2014 following the same protocol described above. As with other attempts to adapt *C. maculatus* to lentil, the line underwent a severe, initial bottleneck. It then persisted at a low census population size (a few hundred beetles per generation with a prolonged period of emergence) for four generations. During that time, there was no evidence of an increase in population size indicative of rescue. We sampled and sequenced 48 adult beetles from the L11 line in the F4 generation. At this point, the culture was discontinued given the lack of evidence of rescue, and thus inability to obtain a sufficient number of beetles for our standard culture procedures. This does not preclude the possibility that the line would have eventually been rescued before true extinction, but there was no evidence rescue was underway.

#### Our GBS approach

We sampled and isolated genomic DNA from 48 adult *C. maculatus* in each of the following generations: L14 founders (P), F1, F2, F3 and F4, L14 subline A (L14A) F5, F6, F7, F8 and F16, L14 subline B (L14B) F5, F8 and F16 (L14A F5 and L14B F5 denote the subsets of the L14 line that founded each of the sublines in the F5 generation), and L11 F4. Adult beetles were collected at the end of each generation, and stored at -80° C. Beetles were surface sterilized by washing in hydrogen peroxide and ethanol prior to extraction. We then isolated and purified DNA from the 672 beetles using Qiagen's DNeasy 96 Blood & Tissue kit (Qiagen Inc.). We generated DNA fragment libraries for genotyping-by-sequencing (GBS) from each beetle following previously published laboratory methods (Gompert *et al.*, 2012), with modifications documented in Gompert *et al.* (2014a). Briefly, we targeted a random subset of the genome by digesting genomic DNA from each beetle with the restriction endonucleases *EcoRI* and *MseI*. We then ligated adaptor oligos with internal 8-10 bp barcode sequences to the sticky ends of the digested DNA using T4 DNA ligase. We PCR amplified the restriction fragment library with the Illumina PCR primers and iProof high-fidelity polymerase (Bio-Rad Laboratories Inc., Hercules, CA, USA). DNA libraries were then combined and size-selected using a BluePippin (Sage Science, Beverly, MA, USA); 250-350 bp fragments were retained. Purified genomic libraries were then sequenced at the University of Texas Genomic Sequencing and Analysis Facility (Austin, TX, USA) on the Illumina HiSeq 4000 platform. Four lanes of 100 bp single-end reads were generated.

#### *De novo* assembly of a *C. maculatus* genome

Heterozygosity can create problems for *de novo* genome assembly. To minimize this issue, we created inbred *C. maculatus* lines using a full-sib mating design. Two lines were used for genome sequencing and assembly, both of which were derived from the base Indian mung bean line (SI). High molecular weight DNA was extracted from pooled samples (≈20 beetles each) from each of these inbred lines by Lucigen Corporation.

A fragment paired-end library and 3 kb and 8 kb mate-pair libraries were created from the DNA from one of the lines. This line had experienced eight generations of full-sib mating and thus had an inbreeding coefficient of $F = 0.826$. A 20 kb mate-pair library was constructed from the pooled DNA of the second line, which had undergone nine generations of full-sib mating and had an inbreeding coefficient of $F = 0.859$. Fragmented DNA was used to construct libraries following Lucigen's protocols for NxSeq DNA Sample Prep Kit (Cat. No. 12000-1) and NxSeq Long Mate Pair Library Kit (Cat. No. 13000-1). The fragment library was sequenced on an Illumina HiSeq 2500 (one lane) with 2×100 bp reads. The 3 kb library and (combined) 8 and 20 kb libraries were each sequenced on a MiSeq with 2×250 bp reads. Library construction, quality controls and sequencing were all performed by Lucigen Corporation.

Sequencing resulted in 315 million reads for the fragment library, 22 million reads from the 3 kb library, 12 million reads from the 8 kb library, and 9 million reads from the 20 kb library. We generated a *de novo* genome assembly from these data using `allpaths-lg` (version 52488) (Gnerre *et al.*, 2011), which was run with `HAPLOIDIFY=False`, `FIX_LOCAL=True` and `MIN_CONTIG=250`.

The assembly resulted in 119,214 scaffolds with a total scaffold length of 939.8 mb. The estimated genome size was 1178.2 mb and thus our assembly comprised about 80% of the estimated genome size (50% of the genome was estimated to be repetitive). The genome assembly was highly fragmented (Fig. S1). The scaffold N50 was 35 kb and there were on average 126.9 scaffolds per mb. We further assessed the completeness and quality of the assembly with `busco` version 1.2. We found 877 complete of the Benchmarking Universal Single-Copy Orthologs (BUSCOs) (32%), 615 fragmented BUSCO's (22%) and were missing 1183 BUSCO's (44%) (this set of genes is expected to be in most/all Eukaryotes). The large number of missing BUSCO's likely reflects the highly fragmented nature of the assembly. Nonetheless, this partial/draft assembly represents a substantial improvement over the best currently available assembly (85,859 scaffolds with a N50 of 3.8 kb, see `https://beanbeetle.org/`; Gompert & Messina, 2016), and provides an improved reference sequence for aligning our re-sequence data and measuring allele frequency change over time.

#### Alignment and variant calling

We used the `aln` and `samse` algorithms from `bwa` (ver. 0.7.10) (Li & Durbin, 2009) to align the 764 million ~86 bp DNA sequences (after trimming barcodes) to a new draft genome assembly for *C. maculatus* (Fig. S1). We allowed a maximum of four missmatches, with no more than two missmatches in the first 20 bps. Sequences were only placed if they had a unique best hit (`-n 1` in `samse`).

We then identified SNPs using the Bayesian multiallelic/rare variant caller from `samtools` (version 1.5) and `bcftools` (version 1.6) (implemented with the `-m` option in `bcftools call`). The `-C 50` command was used, as recommended for Illumina HiSeq data. We ignored bases with a base quality <30 and reads with a mapping quality <20. Variants were only called if the posterior probability that the population was fixed for the reference allele was <0.01 assuming $\Theta = 0.001$.

We then filtered the SNP set to only retain those with minimum coverage of 1344× (across all samples), at least 10 sequences supporting the non-reference allele, a minimum mapping quality of 30, missing data for no more than 201 individuals, a minimum minor allele frequency of at least ~0.005, and no more than 1% of reads in the reverse orientation (with our GBS approach, all reads should be in the same orientation). We applied a second filter to remove reads with a maximum coverage of >21,556× (two SDs greater than the mean) and that were within 4 bp of each other (this pair of filters should remove highly repetitive regions and those that represent poor alignments). 21,342 SNPs were retained following filtering.

#### Allele frequency model

We used a hierarchical Bayesian model to estimate allele frequencies in each (sub)line and generation for each of the 21,342 SNPs, as described in Gompert & Messina (2016). We describe the model again here for completeness. We first used the pre-computed genotype likelihoods from `bcftools`, which denote the relative likelihoods of each genotype given the sequence data and quality scores, to denote the probability of the data conditional on each genotype, that is $\Pr(\mathbf{d_{ijkl}}|g_{ijkl}, \mathbf{q_{ijkl}})$. Here subscripts denote beetle $i$ from (sub)line $j$ sampled from generation $k$ and genotyped (by sequencing) at SNP locus $l$. $g_{ijkl}$ denotes the unknown genotype written as the count of the non-reference allele (0, 1, or 2), $\mathbf{d_{ijkl}}$ is the vector of base calls, and $\mathbf{q_{ijkl}}$ is the corresponding vector of quality scores. The calculation of $\Pr(\mathbf{d_{ijkl}}|g_{ijkl}, \mathbf{q_{ijkl}})$ is described in detail in Li (2011). We then assumed that the prior probability of observing a genotype is a function of the allele frequencies, specifically that $\Pr(g_{ijkl}|p_{jkl}) \sim \text{binomial}(p = p_{jkl}, n = 2)$. $p_{jkl}$ denotes the unknown non-reference allele frequency. Strictly speaking this model assumes that the two alleles at a locus are independent conditional on the allele frequencies, but it serves as reasonable prior as long as the probability of observing an allele in an individual is proportional to the population allele frequency, which should generally be true. We then placed an uninformative beta prior, $p_{jkl} \sim \text{beta}(\alpha = 0.5, \beta = 0.5)$ on the population allele frequencies. This model lacks a closed-form solution, so we instead generate samples from the posterior distribution using MCMC. This was done using a computer program written in `C++` using the Gnu Scientific Library (Gough, 2009), as described in Gompert & Messina (2016).

#### Bayesian bootstrap

We estimated variance effective population size from patterns of allele frequency change across generations using a Bayesian bootstrap method (Jorde & Ryman, 2007; Foll *et al.*, 2015; Gompert, 2016). The specific model used was described fully in Gompert & Messina

(2016). Here we briefly describe the model again for completeness. Allele frequency change between generations was calculated as $Fs = \frac{\sum_t (p_{jk+1l} - p_j kl)^2}{\sum_l (\pi_{jkl}(1-\pi_{jkl}))}$ (Jorde & Ryman, 2007; Gompert, 2016; Gompert & Messina, 2016). An unbiased estimate of $N_e$ was then taken as the mean of $\frac{t}{2Fs'}$, where $t$ is the number of generations between samples and *Fs'* is a correction of *Fs* from eqn. 14 in Jorde & Ryman (2007). The correction assumes a census size of 2000 beetles. 1000 bootstrap replications were performed with a uniform prior placed on *Ne* with bounds placed on 5 and 2000. The medians of the posteriors were used for further analysis and parameterization of subsequent models of evolution by drift. We focus on the median (rather than the mode or mean) of the posterior distribution as it is more closely related to the equal-tail probability intervals (ETPIs) we use to characterize parameter uncertainty (i.e., the median will always be within the 95% ETPIs, but the same need not be true of the mean or mode). *Ne* for lines L1-L3 were calculated between their respective generation and M14.

### Sensitivity to model assumptions

Results in the main text used a SD of 0.3 for the slab component of the mixture prior on the selection coefficients (**s**). Here we show the results of slab standard deviations ranging from 0.1 to 0.5, summarized in Figs. S13–S16. For selection from the parental to F4 generation, a standard deviation of 0.1 produced no credible estimates of selection and a mean point estimate of $|s|$ of 0.022. With a SD of 0.2, we found 12 credible estimates of selection during these early generations, with a mean point estimate of $|s|$ of 0.146. Both the number of credible estimates and the magnitude of selection for point estimates of selection are much lower than those produced with a standard deviation of 0.3 (53 credible interval estimates and mean absolute $s$ of 0.388). The inability to infer credible selection when $N_e$ is low and the prior is constraining $s$ to be close to 0 is not surprising. As expected, higher standard deviations of 0.4 and 0.5 produce more credible estimates than found with a standard deviation of 0.3 (58 and 66, respectively). Even though the number of SNPs with credible effects of selection varied based on the prior SD, point estimates of $s$ were often highly correlated across SDs (Fig. S17). Correlations were weakest with the SD 0.1 and selection during the first four generations ($r \leq 0.33$); the mean for all other comparisons was $\bar{r} = 0.845$ (range = 0.676–0.962). Likewise, choosing different SDs has little effect on the extent to which estimates of $s$ are correlated across sublines and time intervals (Table S2).

We also considered the effect of model averaging on selection coefficient estimates. By including and averaging over constrained models, we could inflate the correlations of $s$ over time or across sublines, particularly if there is little information in the data on which model is best. To assess this, we reran ABC analyses on unconstrained models. Five million new simulations were run with the fully unconstrained model and $s$ was estimated from these using the same methods used previously (see Table S5 and Fig. S18). The numbers and magnitudes of credible $s$ estimates remained similar (unconstrained: P–F4, number = 57, $|\bar{s}| = 0.671$; F4–F16A, number = 44, $|\bar{s}| = 0.392$; F4–F16B, number = 45, $|\bar{s}| = 0.435$) to the results of model averaging (model average: P–F4, number = 53, $|\bar{s}| = 0.746$; F4–F16A, number = 53, $|\bar{s}| = 0.421$; F4–F16B, number = 51, $|\bar{s}| = 0.438$). Whereas the sign of $s$ never changed between sublines or time intervals with our main results, this did occur for one SNP when comparing P–F4 and F4–F16A, and two SNPs when comparing P–F4 and F4–F16B with the unconstrained model (Fig S21). Finally, correlations between sub-lines remain high relative to correlations through time (P–F16A=0.194; P–F16B=0.268; F16A–F16B=0.382), which is consistent with the model averaged results despite lower overall magnitudes.

### Evaluation of the ABC approach

We used simulated output from the ABC simulations above as input to test the ability of our ABC approach to accurately estimate the strength of selection, $s$. We specifically evaluated the method based on the output from 10 simulations based on each of the 198

SNPs (1980 simulated data sets). This was done to ensure we considered a variety of initial allele frequencies as these were based on the founder (P) allele frequency for each SNP. For each of the 1980 pseudo data sets, we used the observed allele frequency trajectory as the observed data, and the output from the other 4,999,999 simulations for the SNP in question to perform ABC inference of $s$ as described above. Posterior estimates of $s$ were based on the 1000 simulations with allele frequency vectors that best matched the allele frequency vector for each of the 1980 pseudo data sets. We then compared the observed and estimated values of $s$ for the pseudo data sets.

When considering all simulations for selection estimates from L14P–L14F4, a significant ($P < 0.05$) correlation of $r = 0.386$ was observed, with a root mean square error (RMSE) of 0.202 (Fig. S4). The proportion of simulated data sets where the true value of $s$ was contained within the 95% ETPIs of the posterior estimate was 0.950. The ABC method underestimated the true value of $s$ in 41.5% of cases, and estimated the correct sign in 59.0% of cases.

Although we are interested in SNPs undergoing strong selection, many estimates of $s$ in our ABC evaluation were based upon observed estimates of $s$ close to zero. Therefore, we considered the efficacy of our model for the subset of cases were the true value of $s$ was $|s| > 0$ (Table S6 and Fig. S5). With larger values of $|s|$, we detected greater correlations between true and estimated values for P–F4. These were significant (i.e., $P < 0.05$) except when using a cut-off of $|s| > 0.8$ (here, $r \geq 0.946$, but too few SNPs meet this and more extreme criteria to test for statistical significance). In contrast, for F4–F16, correlations between true and estimated values of $s$ were lower overall, and less effected by the minimum absolute value of $s$.

### Alternative ABC models

No allele frequency data exists for the true ancestors of the L1, L2, and L3 lines. Instead, a contemporary sample of the mung line from which L1–L3 originated from was taken. At the time the mung line sample was taken, 131, 107, and 104 generations had elapsed since the generation of lines L1, L2, and L3, respectively (fewer generations had passed in these lentil lines because of differences in generation times on mung bean versus lentil). We simulated evolution by drift backwards through time using contemporary estimates of variance $N_e$ for the mung bean line provided in Gompert & Messina (2016). Specifically, for each ABC simulation, change by drift each generation back in time was simulated by sampling $p_{t-1} \sim \text{binomial}(p_t, 2N_e)/N_e$ (as in Gompert & Messina, 2016). The simulated ancestral allele frequency in the mung bean line at the point each lentil line was formed was then used for the forward-in-time simulations of evolution for each lentil line according to the Wright-Fisher model with the appropriate $N_e$ for L1–L3 as described in main text. We assumed constant $s$ in each line, but past work with these lines that focused on evidence of genetic trade-offs in host use considered alternative models (different assumptions affected the absolute magnitude of selection, but had little effect on the relative magnitudes of $s$ across SNPs or on their credibility, which is our main concern here; Gompert & Messina, 2016). The same spike-slab prior for $s$ in the L14 analysis was implemented for lines L1-L3. The modified version of the `C++` software for these simulations is denoted `wfabc-const`. The posterior distribution of $s$ was composed of the closest 0.02% simulations to the last sampled allele frequency for each line (L1-F100, L2-F87, L3-F85) rather than a full allele frequency trajectory as in L14. The `loclinear` method was not used in the L1-L3 line due to a lack of variance in summary statistics. Rather, the default `rejection` method was implemented.

A third version of the ABC model was used for the L11 line. Like the version described in the main text, this version initiated simulations with the observed ancestral allele frequency (we used the founders of L14 for this), and thus backwards-in-time simulations were not necessary. Other than this, this version of the software (dubbed `wfabc-simple`) was identical to `wfabc-const` in that only the last sampled allele frequency for each line (in this case the F4 generation of L11) was used for the ABC analysis.

#### Heterozygous effect

Our ABC model allowed for joint inference of selection coefficients ($s$) and dominance (i.e., the heterozygous effect, $h$). Only two estimates of $h$ had credible intervals which did not span 0.5 in any time interval or subline. Such uncertainty in $h$ is unsurprising and our main reason for including $h$ in the ABC model was to integrate over uncertainty in $h$ instead of assuming perfect additivity.

Nonetheless and despite the overall uncertainty in estiamtes of $h$, we detected consistent and significant but weak correlations between point estimates of $s$ and $h$ (L14 P–F4, $r_{s,h} = -0.107$; L14A F4–F16, $r_{s,h} = -0.048$; L14B F4–F16, $r_{s,h} = -0.045$; Table S3, Fig. S20). Estimates of $s$ and $h$ were most strongly correlated early in the adaptation processes. These results are consistent with a general trend of selection favoring initially rare, recessive alleles. Additional work to further test this hypothesis is warranted.

#### Supplemental Tables and Figures

| Generation | 1-100 bp | 101-1000 bp | 1001 bp–10 kbp | 10–500 kbp | Different scaffolds |
|---|---|---|---|---|---|
| LD (mean) | | | | | |
| P | 0.369 | 0.157 | 0.173 | 0.152 | 0.026 |
| F1 | 0.367 | 0.148 | 0.157 | 0.142 | 0.025 |
| F4 | 0.382 | 0.152 | 0.163 | 0.147 | 0.032 |
| F16A | 0.419 | 0.071 | 0.069 | 0.066 | 0.043 |
| F16B | 0.428 | 0.080 | 0.083 | 0.078 | 0.052 |
| LD (median) | | | | | |
| P | 0.126 | 0.036 | 0.046 | 0.041 | 0.011 |
| F1 | 0.124 | 0.035 | 0.043 | 0.040 | 0.011 |
| F4 | 0.123 | 0.034 | 0.037 | 0.035 | 0.010 |
| F16A | 0.160 | 0.029 | 0.031 | 0.027 | 0.017 |
| F16B | 0.190 | 0.033 | 0.034 | 0.032 | 0.021 |
| LD (90th percentile) | | | | | |
| P | 0.998 | 0.578 | 0.579 | 0.514 | 0.067 |
| F1 | 0.998 | 0.500 | 0.506 | 0.475 | 0.064 |
| F4 | 0.998 | 0.497 | 0.524 | 0.487 | 0.077 |
| F16A | 0.995 | 0.201 | 0.188 | 0.186 | 0.117 |
| F16B | 0.995 | 0.215 | 0.224 | 0.215 | 0.143 |

Table S1: Mean, median, and 90th percentile LD ($r^2$) for SNPs within different bp distances of each other which share scaffolds, and for SNPs on different scaffolds.

| Lines | $r$ (median) | $r$ (95% ETPIs) |
|---|---|---|
| SD=0.1 | | |
| L14F4–L14F16A | 0.440 | 0.224–0.607 |
| L14F4–L14F16B | 0.446 | 0.227–0.600 |
| L14F16A–L14F16B | 0.842 | 0.681–0.911 |
| SD=0.2 | | |
| L14F4–L14F16A | 0.518 | 0.375–0.633 |
| L14F4–L14F16B | 0.528 | 0.404–0.635 |
| L14F16A–L14F16B | 0.861 | 0.772–0.910 |
| SD=0.3 | | |
| L14F4–L14F16A | 0.489 | 0.373–0.587 |
| L14F4–L14F16B | 0.499 | 0.387–0.592 |
| L14F16A–L14F16B | 0.857 | 0.753–0.914 |
| SD=0.4 | | |
| L14F4–L14F16A | 0.451 | 0.304–0.563 |
| L14F4–L14F16B | 0.469 | 0.329–0.587 |
| L14F16A–L14F16B | 0.825 | 0.689–0.897 |
| SD=0.5 | | |
| L14F4–L14F16A | 0.468 | 0.334–0.570 |
| L14F4–L14F16B | 0.485 | 0.354–0.587 |
| L14F16A–L14F16B | 0.817 | 0.680–0.887 |

Table S2: Pearson correlations ($r$) between estimates of selection ($s$) for different sublines and time intervals using different prior SDs for the slab component of the prior on $s$. Results in the main text use SD = 0.3. Posterior distributions of the Pearson correlation coefficient ($r$) are shown that incorporate uncertainty in estimates of $s$.

| | $r$ (median) | $r$ (95% ETPIs) |
|---|---|---|
| L14P–L14F4 | -0.178 | -0.306– -0.048 |
| L14F4–L14F16A | -0.067 | -0.194–0.062 |
| L14F4–L14F16B | -0.074 | -0.203–0.055 |

Table S3: Pearson correlations ($r$) between estimates of selection ($s$) and dominance ($h$) for different sublines and time intervals. Posterior distributions of the Pearson correlation coefficient ($r$) incorporate uncertainty in estimates of $s$.

| (Sub)line/interval 1 | (Sub)line/interval 2 | $r$ (median) | $r$ (95% ETPIs) |
|---|---|---|---|
| L1 | L2 | 0.115 | 0.001–0.226 |
| L1 | L3 | 0.094 | -0.032–0.235 |
| L2 | L3 | 0.262 | 0.139–0.393 |
| L1 | L14–P to L14–F4 | 0.005 | -0.095–0.118 |
| L2 | L14–P to L14–F4 | -0.027 | -0.131–0.081 |
| L3 | L14–P to L14–F4 | -0.015 | -0.136–0.103 |
| L1 | L14–F4 to L14A–F16 | 0.048 | -0.066–0.150 |
| L2 | L14–F4 to L14A–F16 | -0.024 | -0.159–0.102 |
| L3 | L14–F4 to L14A–F16 | -0.006 | -0.133–0.117 |
| L1 | L14–F4 to L14B–F16 | 0.050 | -0.061–0.151 |
| L2 | L14–F4 to L14B–F16 | -0.019 | -0.151–0.095 |
| L3 | L14–F4 to L14B–F16 | -0.019 | -0.145–0.099 |
| L14–P to L14–F4 | L14–F4 to L14A–F16 | 0.489 | 0.373–0.587 |
| L14–P to L14–F4 | L14–F4 to L14B–F16 | 0.499 | 0.387–0.592 |
| L14–F4 to L14A–F16 | L14–F4 to L14A–F16 | 0.857 | 0.753–0.914 |
| L11 | L1 | -0.011 | -0.143–0.120 |
| L11 | L2 | -0.018 | -0.151–0.138 |
| L11 | L3 | -0.032 | -0.196–0.145 |
| L11 | L14–P to L14–F4 | 0.023 | -0.108–0.153 |
| L11 | L14–4 to L14–F16A | 0.003 | -0.121–0.132 |
| L11 | L14–4 to L14–F16B | 0.016 | -0.120–0.155 |

Table S4: Pearson correlations ($r$) between estimates of selection ($s$) for different lines, sublines and time intervals. Posterior distributions of the Pearson correlation coefficient ($r$) incorporate uncertainty in estimates of $s$.

| Subline/interval 1 | Subine/interval 2 | $r$ (median) | $r$ (95% ETPIs) |
|---|---|---|---|
| L14–P to L14–F4 | L14–F4 to L14A–F16 | 0.194 | 0.079–0.305 |
| L14–P to L14–F4 | L14–F4 to L14B–F16 | 0.268 | 0.162–0.365 |
| L14–F4 to L14A–F16 | L14–F4 to L14B–F16 | 0.382 | 0.255–0.504 |

Table S5: Pearson correlations ($r$) between estimates of selection ($s$) for different sublines and time intervals using an **unconstrained model** that posits different values of $s$ for each subline/time interval. Posterior distributions of the Pearson correlation coefficient ($r$) incorporate uncertainty in estimates of $s$.

| | $\lvert s\rvert > 0$ | $\lvert s\rvert > 0.1$ | $\lvert s\rvert > 0.2$ | $\lvert s\rvert > 0.3$ | $\lvert s\rvert > 0.4$ | $\lvert s\rvert > 0.5$ | $\lvert s\rvert > 0.6$ | $\lvert s\rvert > 0.7$ | $\lvert s\rvert > 0.8$ | $\lvert s\rvert > 0.9$ | $\lvert s\rvert > 1$ |
|---|---|---|---|---|---|---|---|---|---|---|---|
| L14P–L14F4 | | | | | | | | | | | |
| Correlation | **0.386** | **0.488** | **0.507** | **0.530** | **0.539** | **0.595** | **0.654** | **0.722** | **0.946** | 1.000 | 1.000 |
| RMSE | 0.202 | 0.319 | 0.368 | 0.428 | 0.496 | 0.564 | 0.648 | 0.739 | 0.809 | 1.091 | 1.091 |
| Underestimate | 0.415 | 0.023 | 0.018 | 0.015 | 0.021 | 0.021 | 0.000 | 0.000 | 0.000 | 0.000 | 0.000 |
| Correct sign | 0.590 | 0.714 | 0.722 | 0.771 | 0.771 | 0.814 | 0.816 | 0.800 | 0.800 | 1.000 | 1.000 |
| Within 95% ETPI | 0.950 | 0.950 | 0.952 | 0.959 | 0.966 | 0.978 | 0.985 | 0.991 | 0.997 | 0.999 | 0.999 |
| L14F4–L14F16A | | | | | | | | | | | |
| Correlation | **0.243** | **0.339** | **0.373** | **0.407** | **0.407** | **0.412** | **0.334** | 0.179 | -0.202 | 1.000 | 1.000 |
| RMSE | 0.241 | 0.345 | 0.388 | 0.445 | 0.514 | 0.590 | 0.690 | 0.792 | 0.949 | 0.876 | 0.876 |
| Underestimate | 0.512 | 0.153 | 0.112 | 0.074 | 0.037 | 0.031 | 0.000 | 0.000 | 0.000 | 0.000 | 0.000 |
| Correct Sign | 0.574 | 0.675 | 0.679 | 0.697 | 0.686 | 0.701 | 0.735 | 0.680 | 0.600 | 1.000 | 1.000 |
| Within 95% ETPI | 0.772 | 0.847 | 0.871 | 0.911 | 0.941 | 0.968 | 0.982 | 0.989 | 0.996 | 0.999 | 0.999 |
| L14F4–L14F16B | | | | | | | | | | | |
| Correlation | **0.218** | **0.296** | **0.323** | **0.341** | **0.350** | **0.313** | 0.281 | -0.071 | -0.246 | 1.000 | 1.000 |
| RMSE | 0.242 | 0.353 | 0.398 | 0.460 | 0.527 | 0.614 | 0.699 | 0.823 | 0.965 | 0.882 | 0.882 |
| Underestimate | 0.501 | 0.135 | 0.102 | 0.074 | 0.043 | 0.041 | 0.000 | 0.000 | 0.000 | 0.000 | 0.000 |
| Correct sign | 0.570 | 0.656 | 0.650 | 0.659 | 0.660 | 0.660 | 0.694 | 0.600 | 0.500 | 1.000 | 1.000 |
| Within 95% ETPI | 0.788 | 0.852 | 0.877 | 0.915 | 0.943 | 0.967 | 0.982 | 0.990 | 0.997 | 0.999 | 0.999 |

Table S6: A summary of metrics used to evaluate the efficacy of our ABC model. Correlation = Pearson correlation coefficients, and bold font denotes values significantly different than zero (i.e., $P < 0.05$). Underestimate = the proportion of estimated $s$ values that were smaller in magnitude than the observed $s$, Correct sign = the proportion of estimated $s$ values that had the same sign as observed $s$

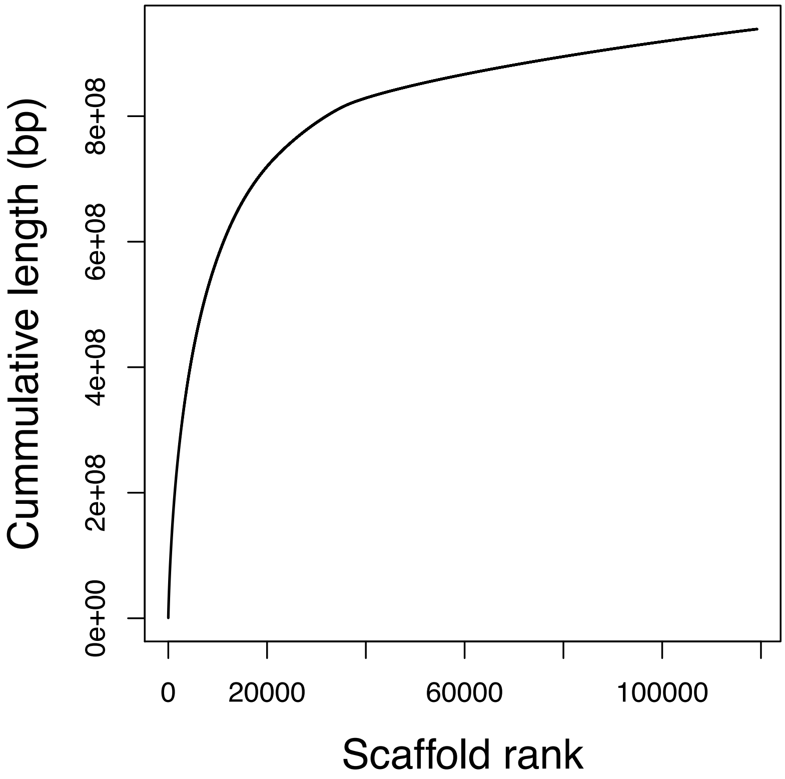


Figure S1: Summary of scaffold sizes. This plots shows the cumulative (summed) genome size for scaffolds sorted by size (from largest to smallest).

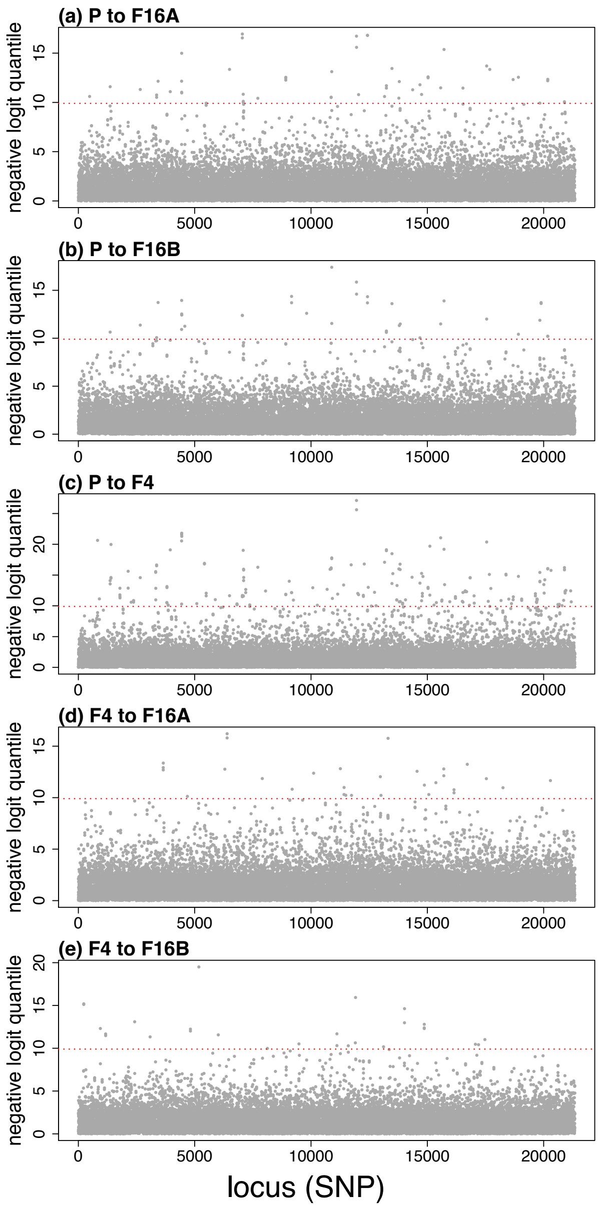


Figure S2: Plots of the percentiles of the null distributions where allele frequency differences fall. The percentiles are logit transformed to more easily observe extreme values. The red line indicates the 0.1th and 99.9th percentiles of the null distribution.

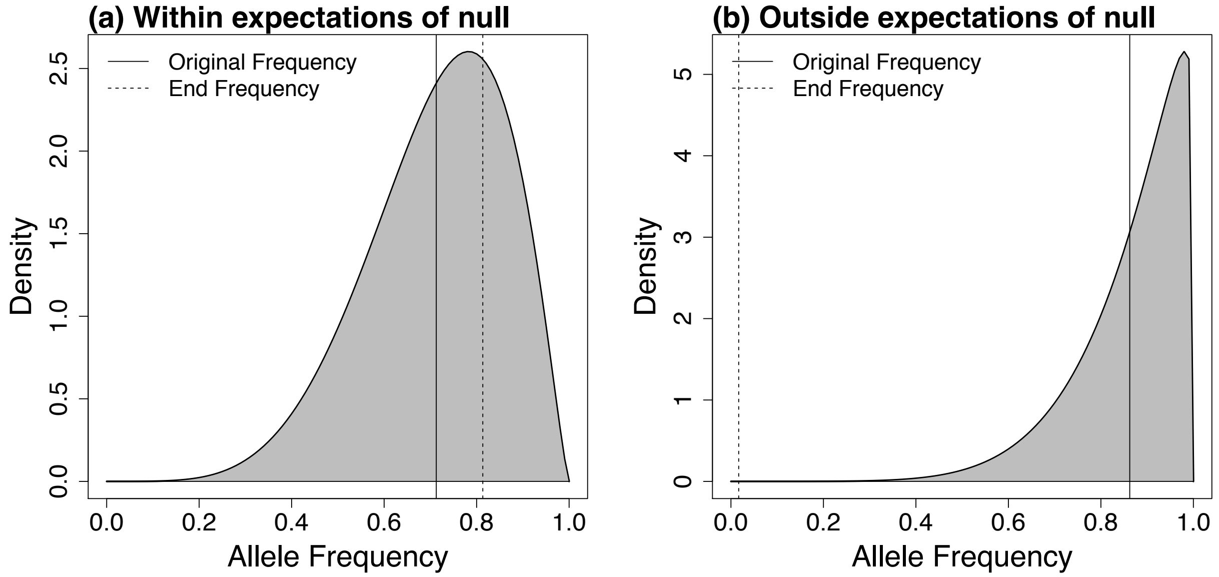


Figure S3: Visual representations of the null distributions for expected change caused by drift. (a) A change in allele frequency that can be easily explained by genetic drift. (b) A change in allele frequency that is unlikely given the null model of genetic drift.

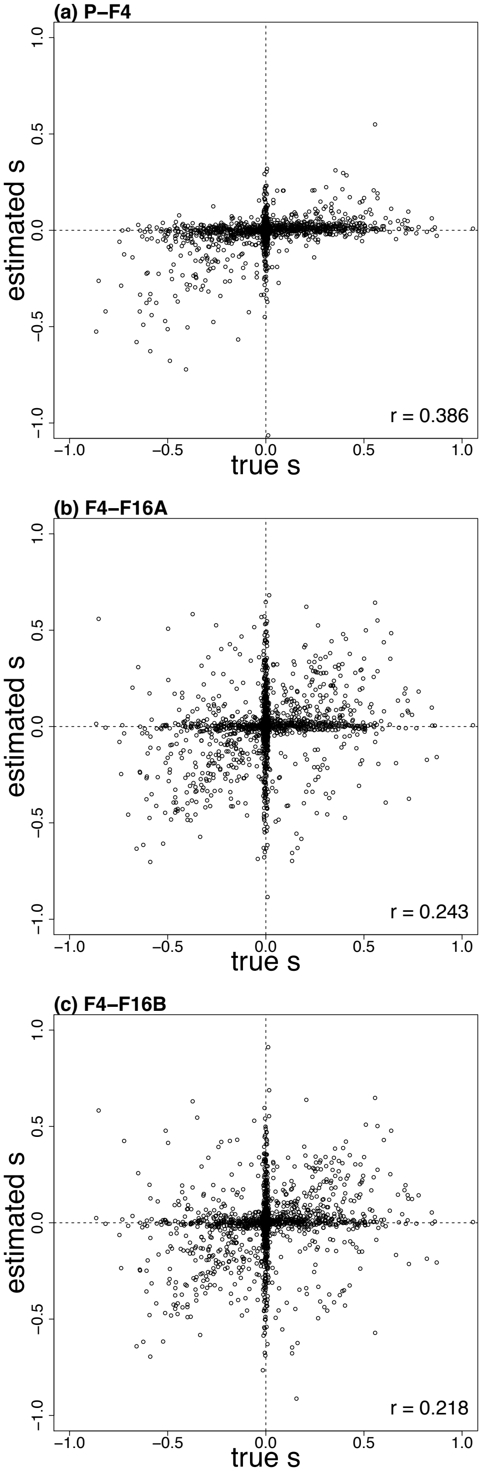


Figure S4: Plot of the ability of the ABC model to estimate $s$ given a known value of $s$. The correlation presented is the simple Pearson correlation coefficient.

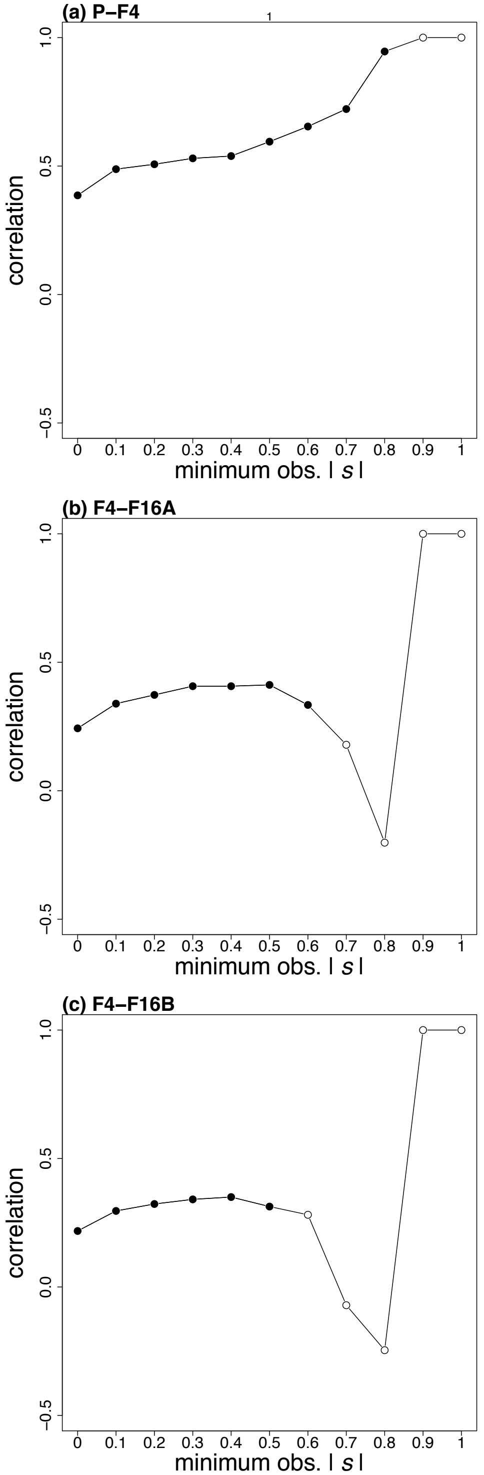


Figure S5: Plots of correlations across thresholds for the minimum true $|s|$ for evaluation the ABC inference procedure. Solid dots denote correlation coefficients that were significantly different from zero ($P < 0.05$), whereas open dots denote correlation coefficients that were not significantly different from zero.

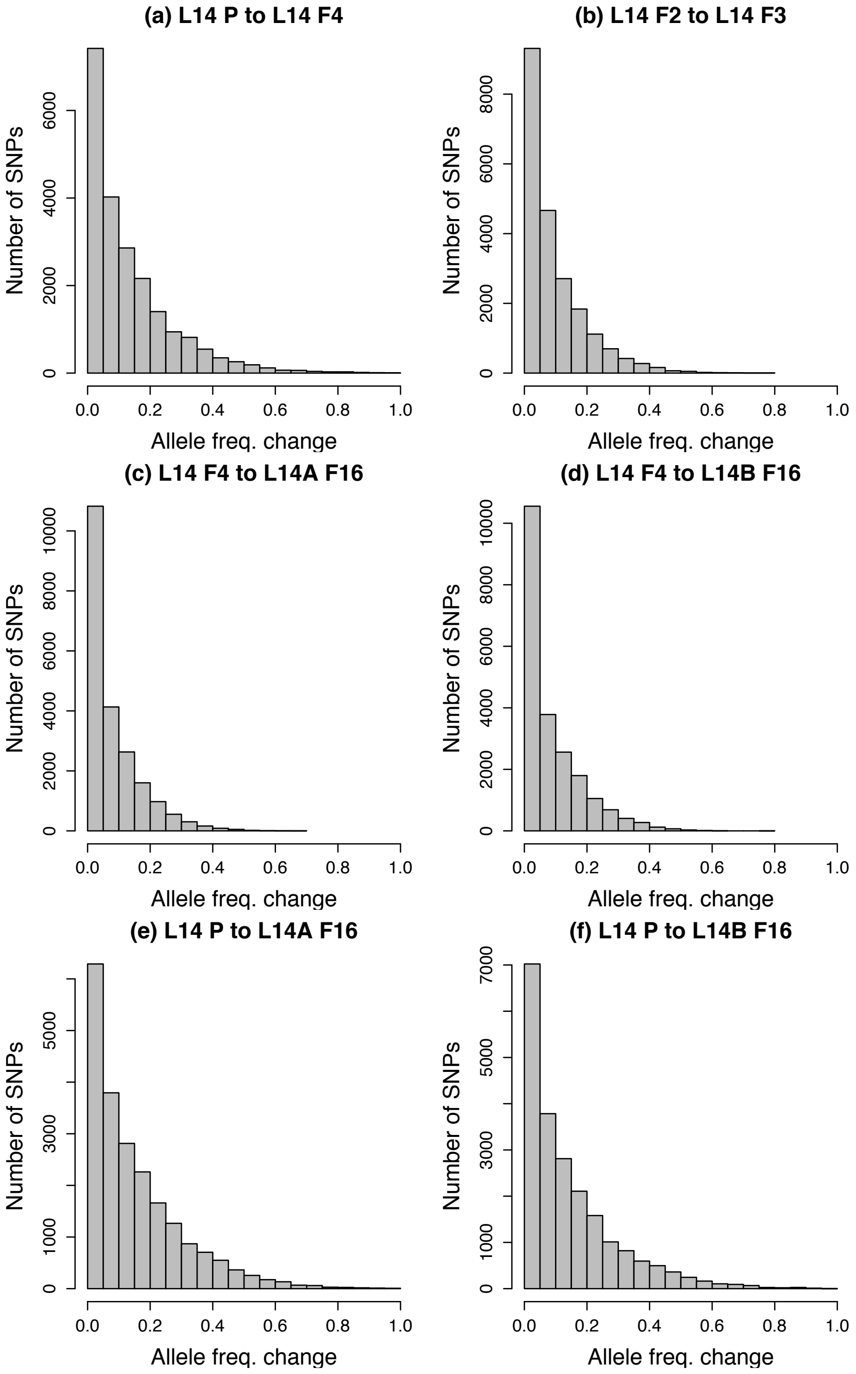


Figure S6: Histograms summarize the distribution of allele frequency change across SNPs (based on the absolute value of allele frequency change). Results are shown for different time intervals and sublines. Some time intervals cover a single generation (e.g., panel b), whereas others span the entire experiment (panels e and f).

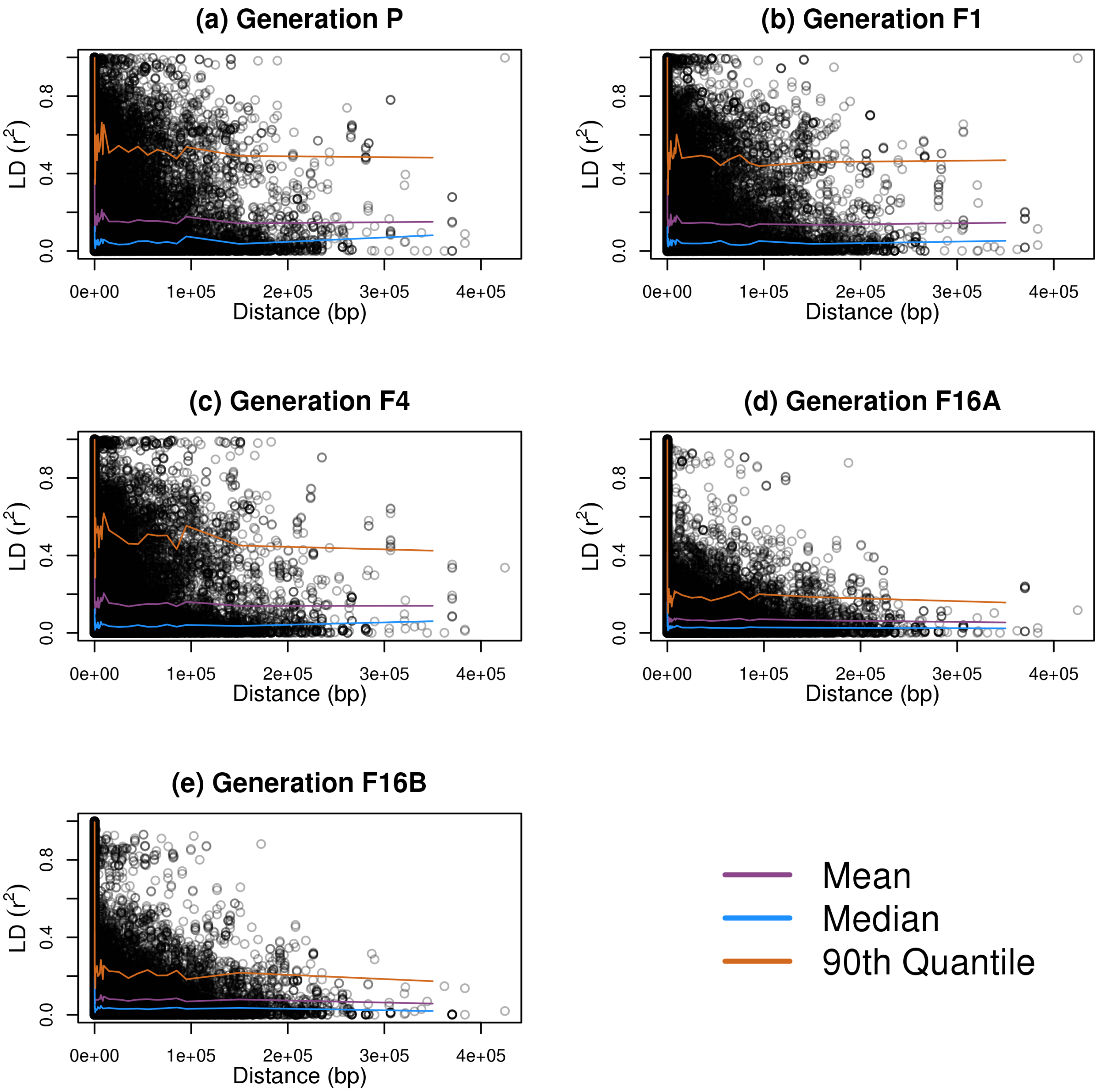


Figure S7: Scatterplots show LD ($r^2$) between pairs of SNPs on the same genome scaffolds as a function of physical distance. Colored lines denote the mean, median and 90th percentile defined over 100, 1000, 10,000, 100,000 or 300,000 bp windows. Results are shown for L14–P (a), L14–F1 (b), L14–F4 (c), L14A–F16 (d) and L14B–F16 (e).

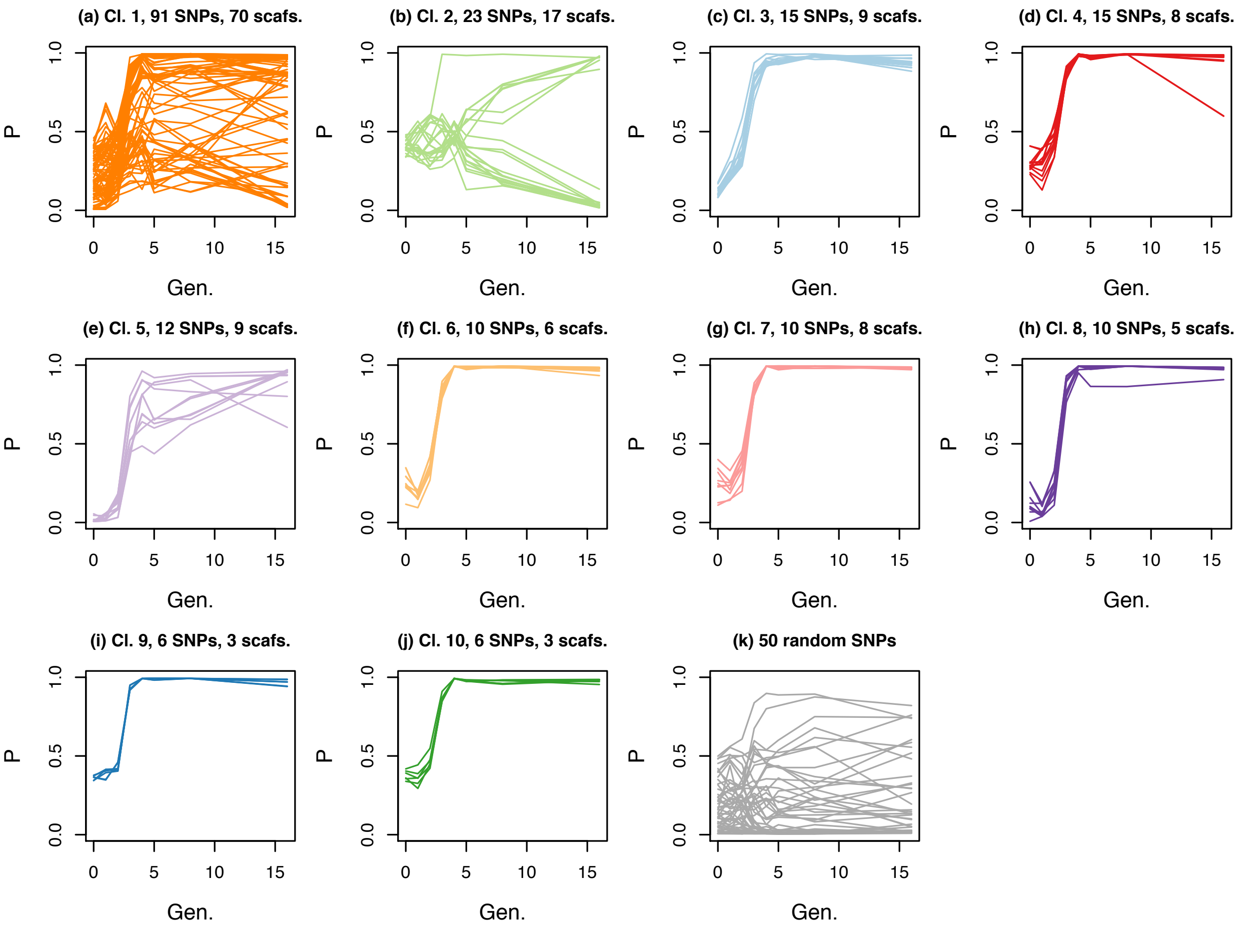


Figure S8: Plots depict patterns of allele frequency change for **L14 subline B**. Panels (a)–(j) show allele frequency ($P$) over time (Gen. = generation) for the 198 focal SNPs. Each line shows the allele frequency trajectory for a single SNP and these are organized into panels by the LD clusters delineated in the F1 generation (Cl. = cluster number; see Fig. 4 and the main text for details). Colors correspond with those from L14–F1 in Fig. 4(a). The number of SNPs and number in each panel and number of scaffolds on which they reside is given. Panel (k) shows patterns of change for 50 randomly selected SNPs. In all cases, the frequency of the minor allele from the parental generation is shown. See Fig. 3 for similar results from L14A.

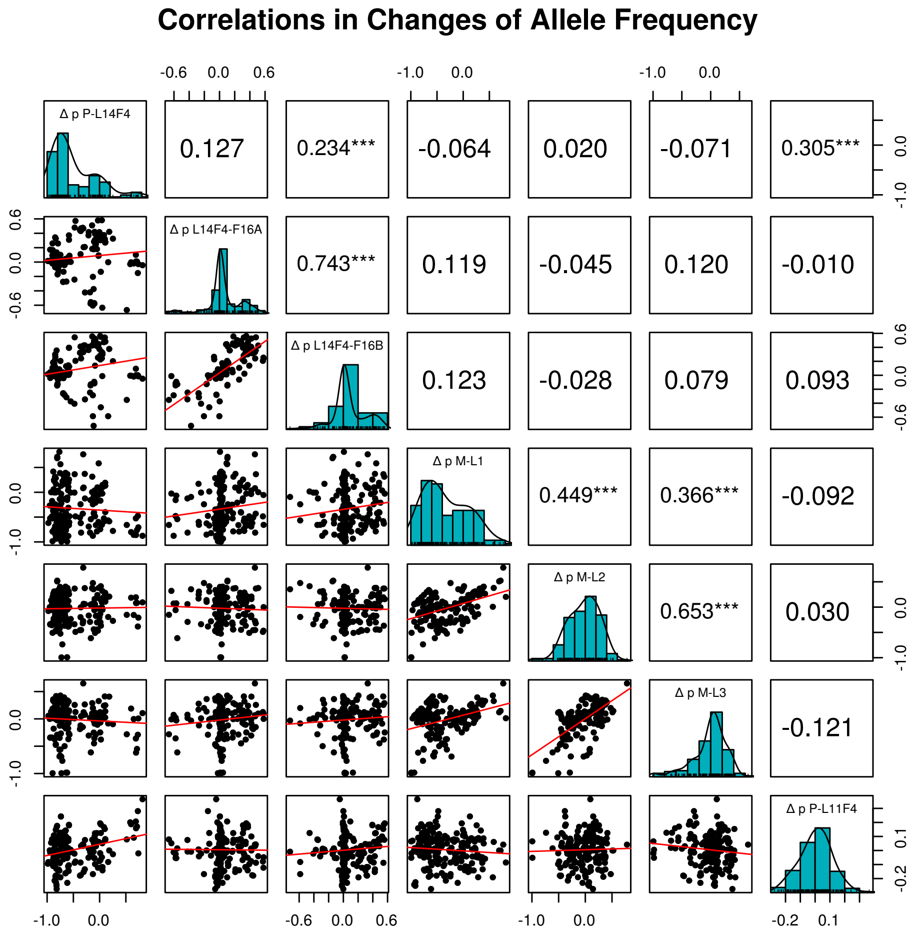


Figure S9: Pairwise comparisons among (sub)lines and intervals of allele frequency change for the 198 focal loci. Comparisons with lines L1, L2, and L3 use only the 188 SNPs present in those lines. The diagonal show the distribution of allele frequency change for an interval. The lower triangle consists of scatter plots of allele frequency change between intervals, and the upper triangle shows correlations between these intervals, with *** denoting significant correlations (i.e., $P < 0.05$).

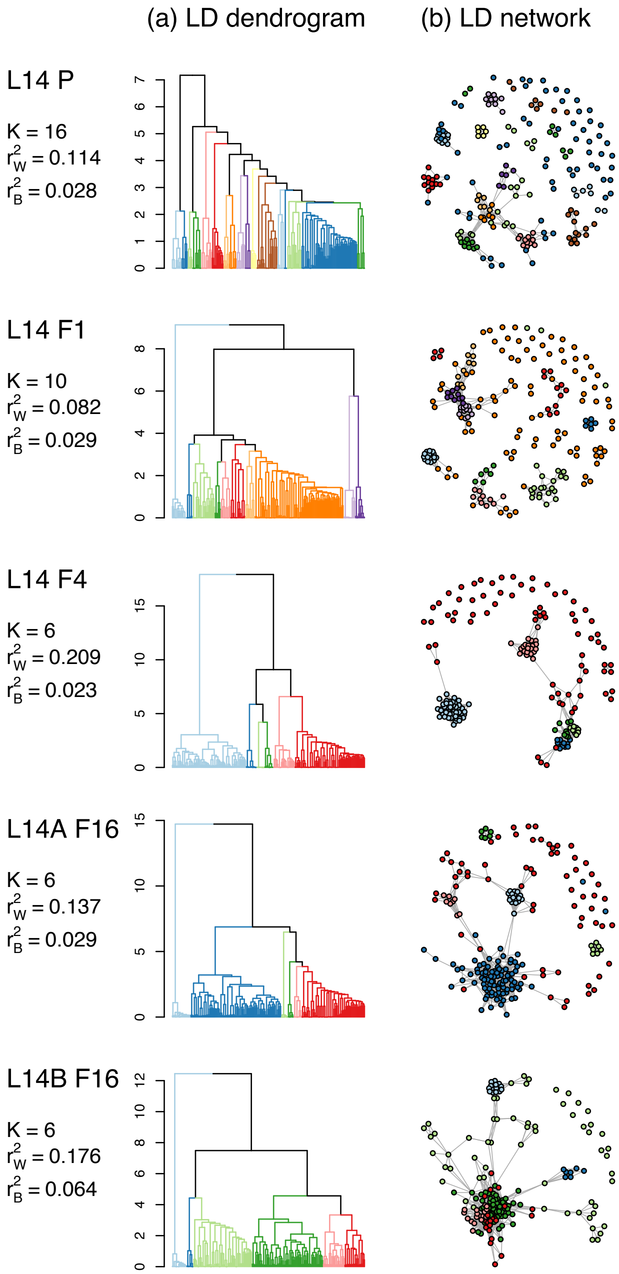


Figure S10: Patterns of LD among the 198 focal SNPs for L14–P, L14–F1, L14–F4, L14A–F16 and L14B–F16. Panel (a) shows dendrograms from hiearchical clustering of SNPs based on LD, with colors denoting clusters delineated with the `cutreeDynamic` function (colors do not track clusters across generations). The number of clusters ($K$) and mean LD for SNPs in the same ($r^2_W$) versus different ($r^2_B$) clusters are given. Panel (b) shows networks connecting SNPs (nodes = colored dots) with high LD ($r^2 \geq 0.25$). Nodes are colored based on their cluster membership from hierachical clustering in the corresponding generation (the same networks are shown but colored based on the F1 generation in Fig. 4).

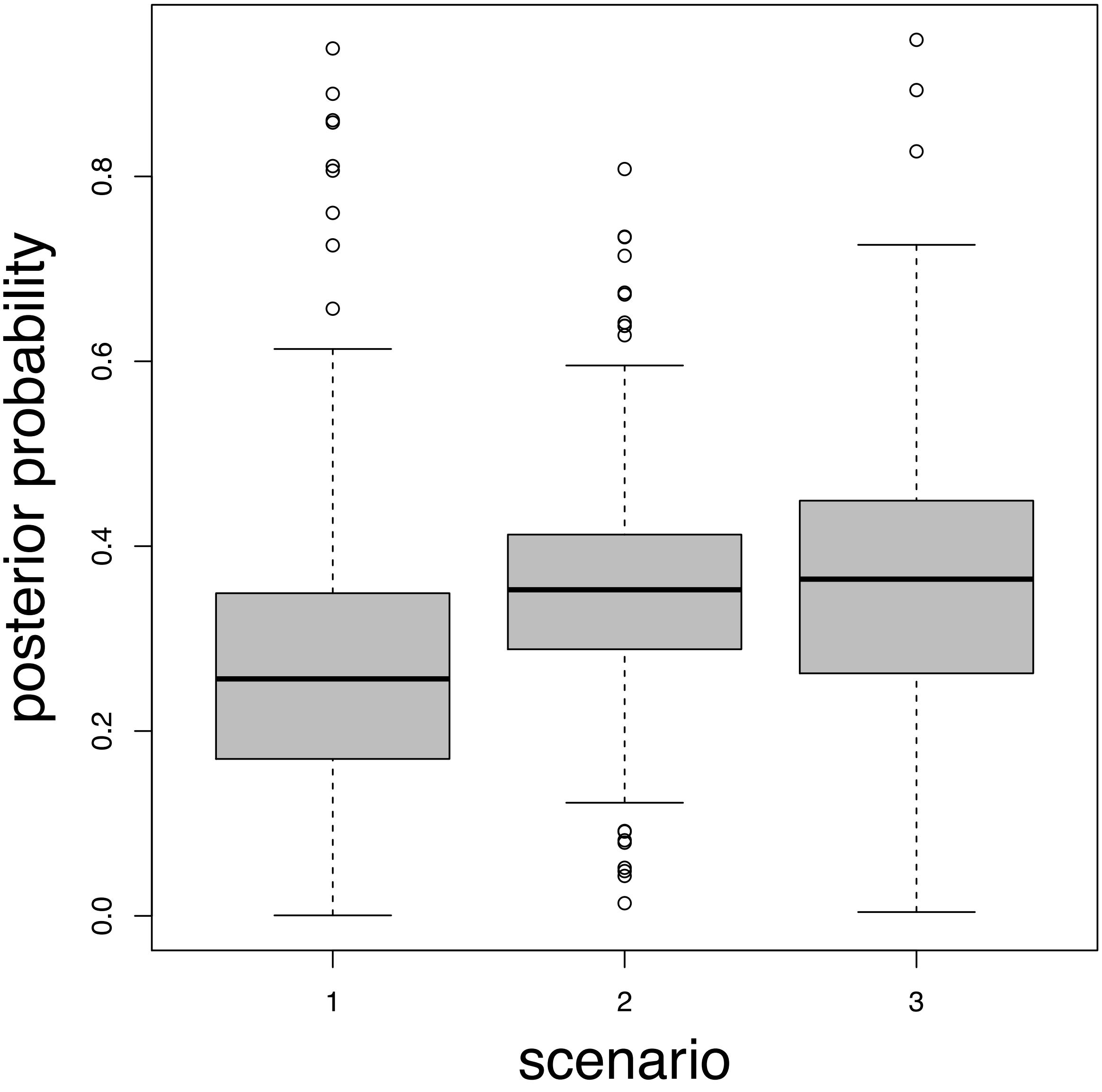


Figure S11: The posterior probability across all loci of each WFABC scenario. Scenario 1 represents a fully constrained model which is not allowed to vary through time or between sub-lines. Scenario 2 represents a model which is allowed to vary through time but not between sub-lines. Scenario 3 represents a fully unconstrained model in which the model is allowed to vary through time and between sub-lines.

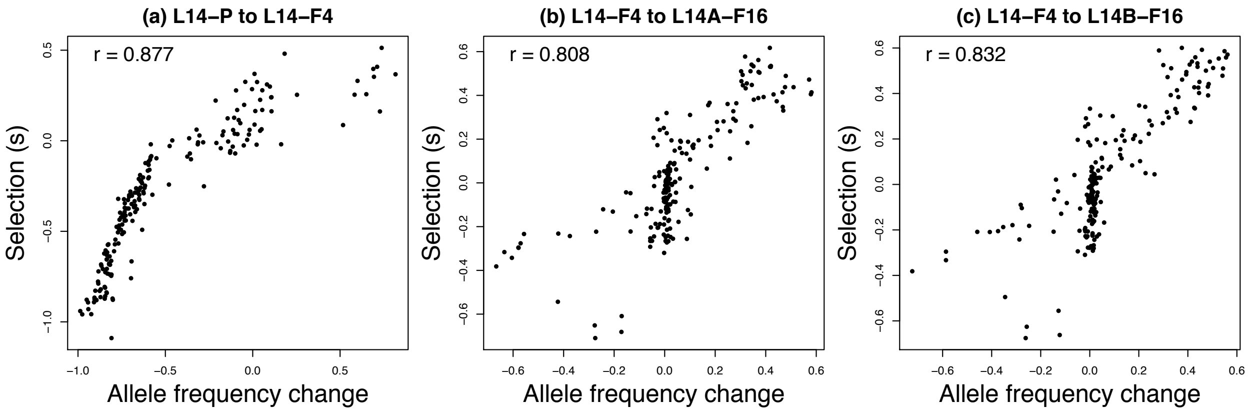


Figure S12: Scatter plots show the relationship between observed allele frequency change and Bayesian selection coefficient estimates ($s$) for different sublines and time intervals for the 198 focal SNPs. Plots are based on point estimates (posterior median) of $s$. Pearson correlation coefficients are shown for each panel.

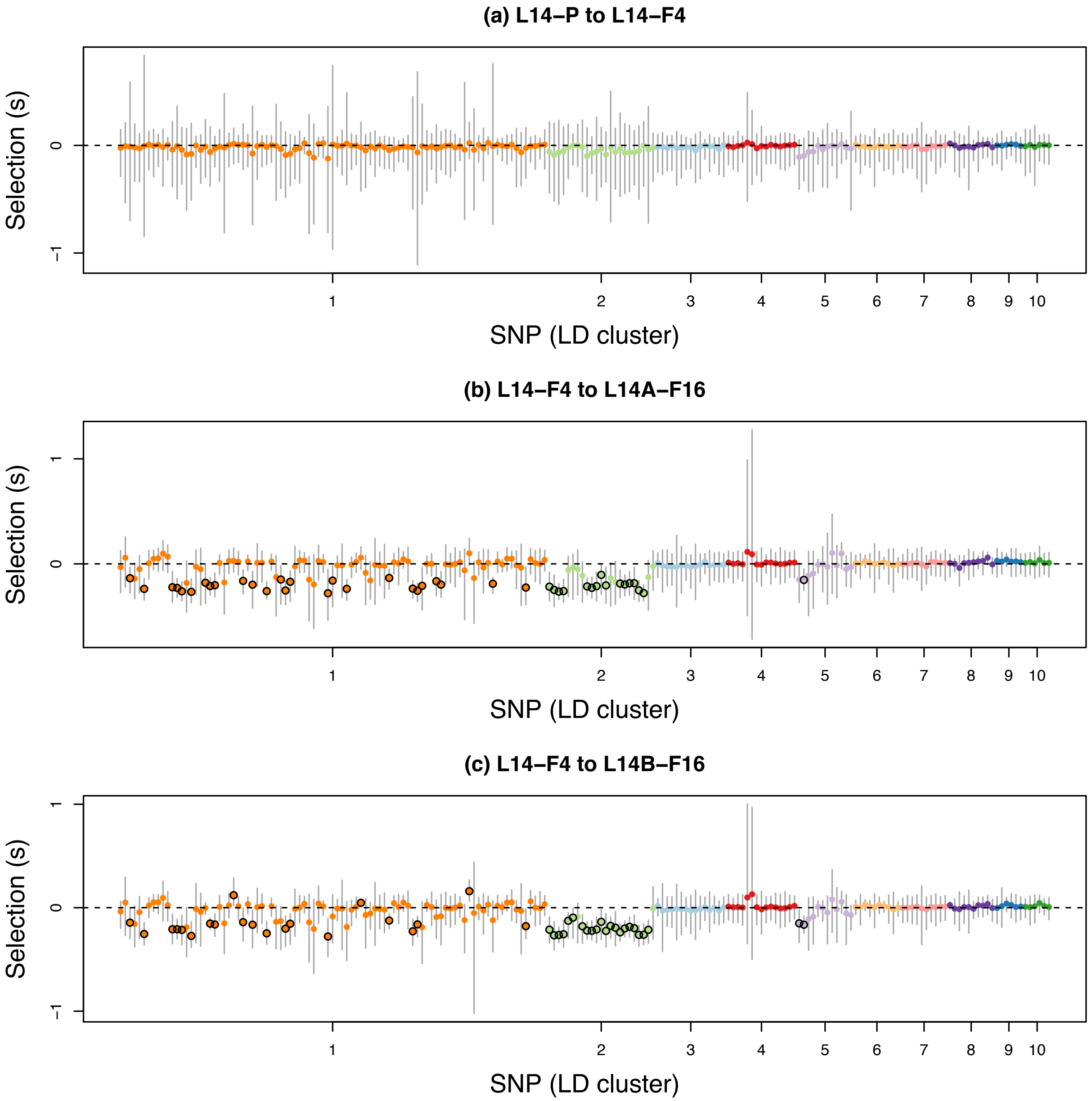


Figure S13: Scatter plots show Bayesian estimates of selection coefficients for the 198 focal SNPs in different generations and sublines with a **prior standard deviation 0.1** for the slab component of the prior on $s$. Dots and vertical bars denote posterior medians and 95% equal-tail probability intervals (ETPIs), respectively. Colors and the order of SNPs reflect LD cluster membership in the F1 generation. Black circles around dots denote cases where the 95% ETPIs exclude 0. For the purpose of visualization, we have polarized estimates of $s$ such that negative values indicate selection favoring the minor allele.

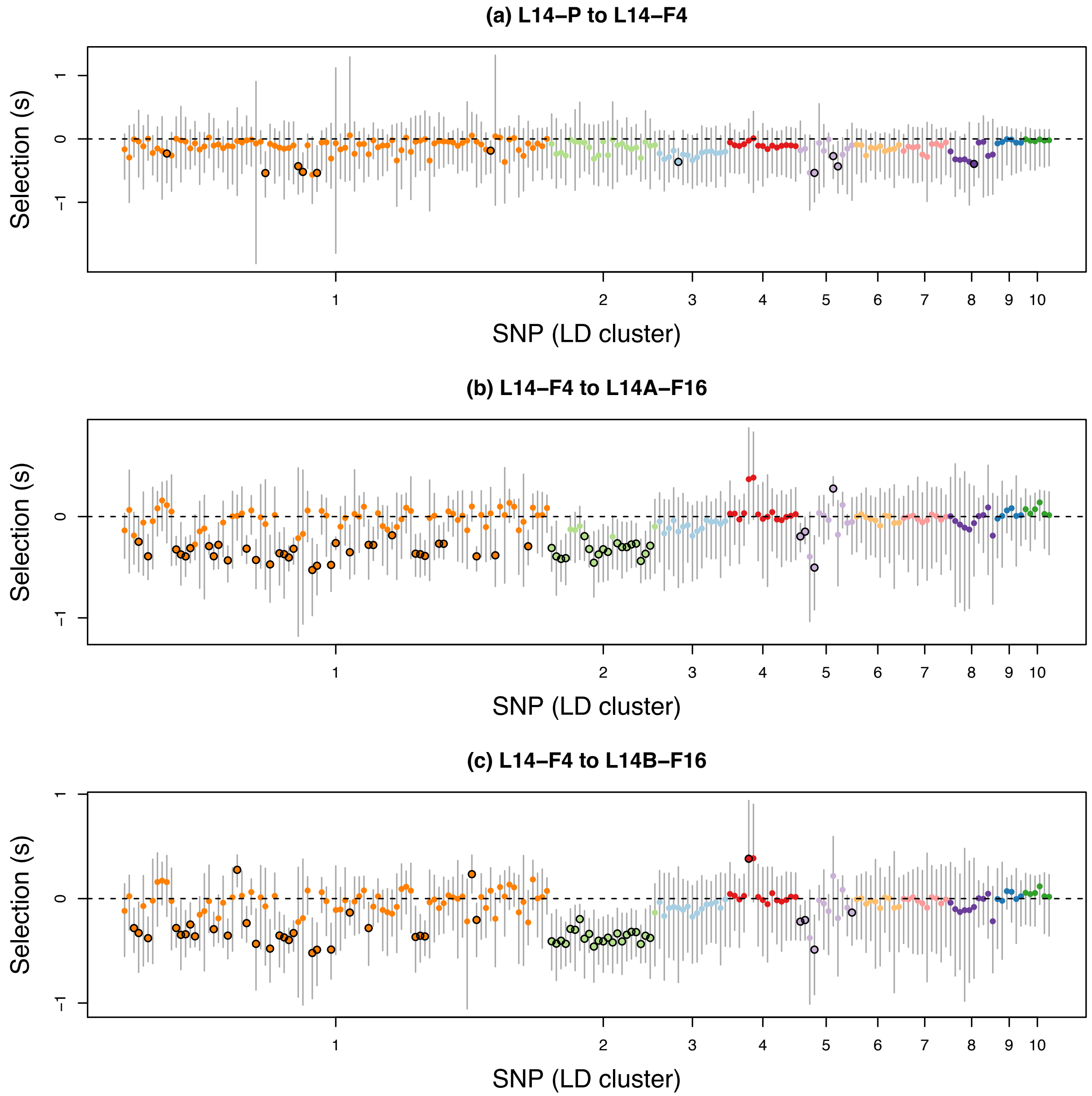


Figure S14: Scatter plots show Bayesian estimates of selection coefficients for the 198 focal SNPs in different generations and sublines with a **prior standard deviation 0.2** for the slab component of the prior on $s$. Dots and vertical bars denote posterior medians and 95% equal-tail probability intervals (ETPIs), respectively. Colors and the order of SNPs reflect LD cluster membership in the F1 generation. Black circles around dots denote cases where the 95% ETPIs exclude 0. For the purpose of visualization, we have polarized estimates of $s$ such that negative values indicate selection favoring the minor allele.

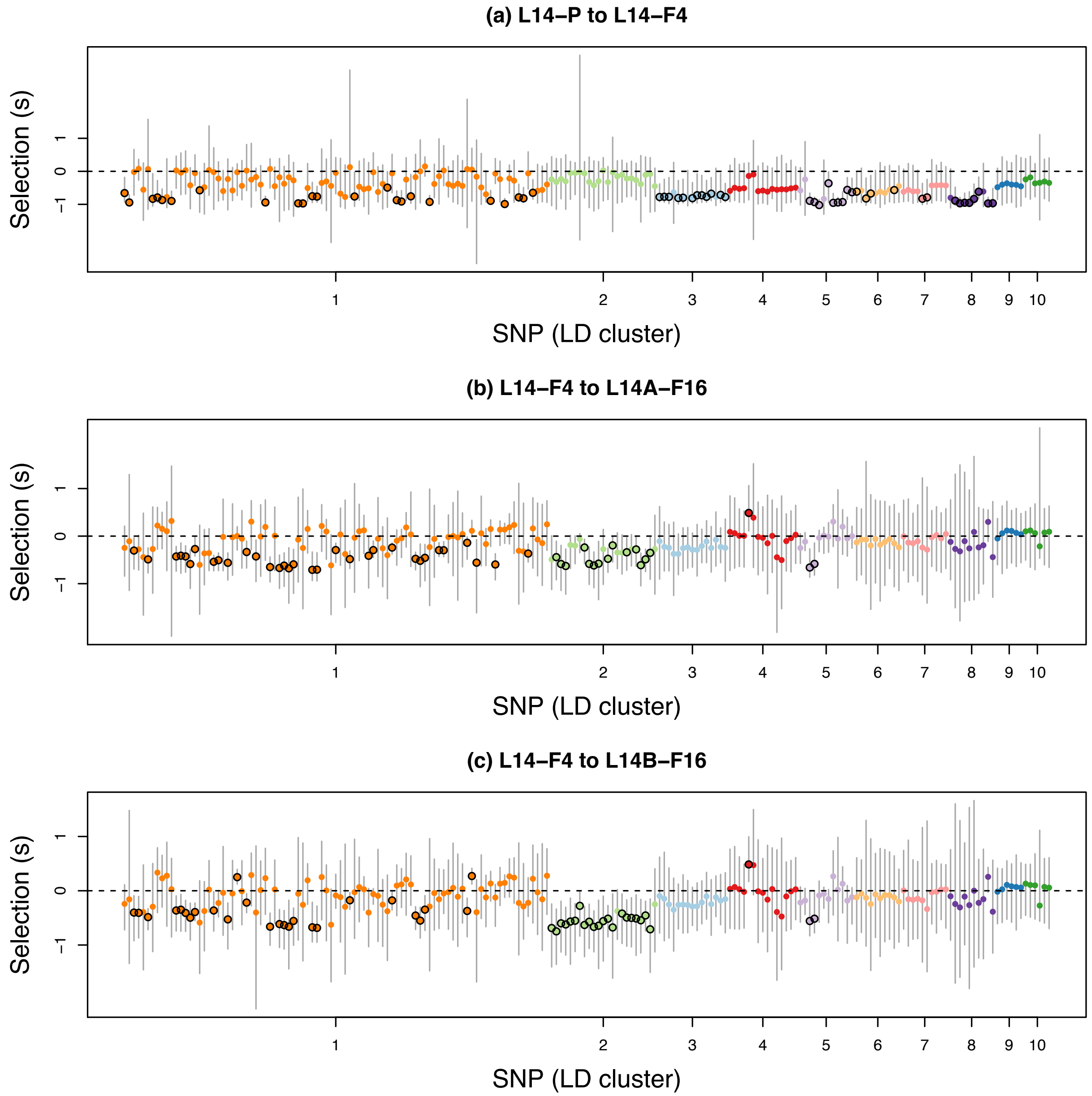


Figure S15: Scatter plots show Bayesian estimates of selection coefficients for the 198 focal SNPs in different generations and sublines with a **prior standard deviation 0.4** for the slab component of the prior on $s$. Dots and vertical bars denote posterior medians and 95% equal-tail probability intervals (ETPIs), respectively. Colors and the order of SNPs reflect LD cluster membership in the F1 generation. Black circles around dots denote cases where the 95% ETPIs exclude 0. For the purpose of visualization, we have polarized estimates of $s$ such that negative values indicate selection favoring the minor allele.

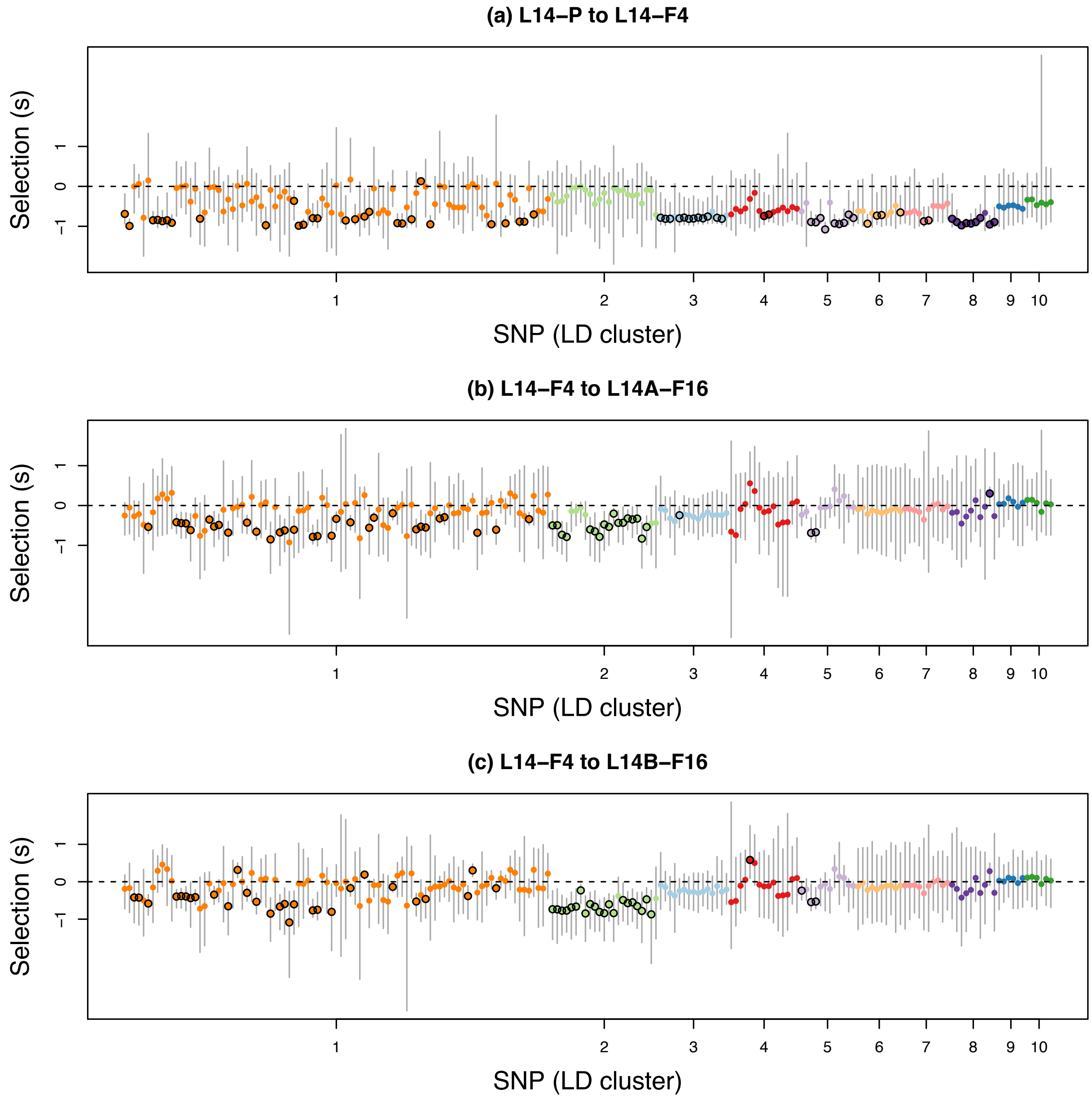


Figure S16: Scatter plots show Bayesian estimates of selection coefficients for the 198 focal SNPs in different generations and sublines with a **prior standard deviation 0.5** for the slab component of the prior on $s$. Dots and vertical bars denote posterior medians and 95% equal-tail probability intervals (ETPIs), respectively. Colors and the order of SNPs reflect LD cluster membership in the F1 generation. Black circles around dots denote cases where the 95% ETPIs exclude 0. For the purpose of visualization, we have polarized estimates of $s$ such that negative values indicate selection favoring the minor allele.

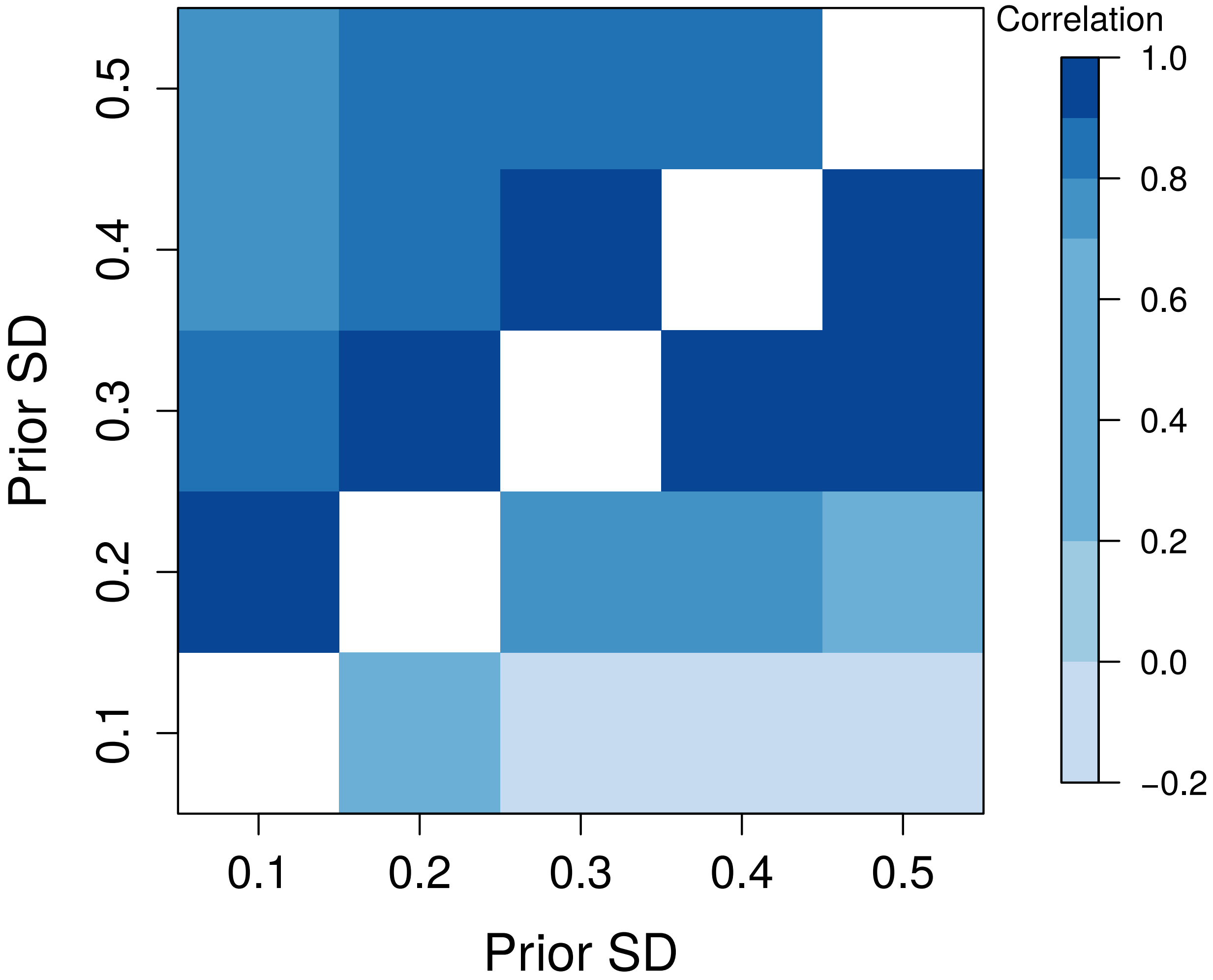


Figure S17: Heat map summarizing the Pearson correlations between selection coefficient point estimates with different SDs for the slab component of the prior. Colored boxes denote correlations coefficients. The upper triangle shows results for selection in subline A after the split, whereas the lower selection shows results for selection from generation P to F4.

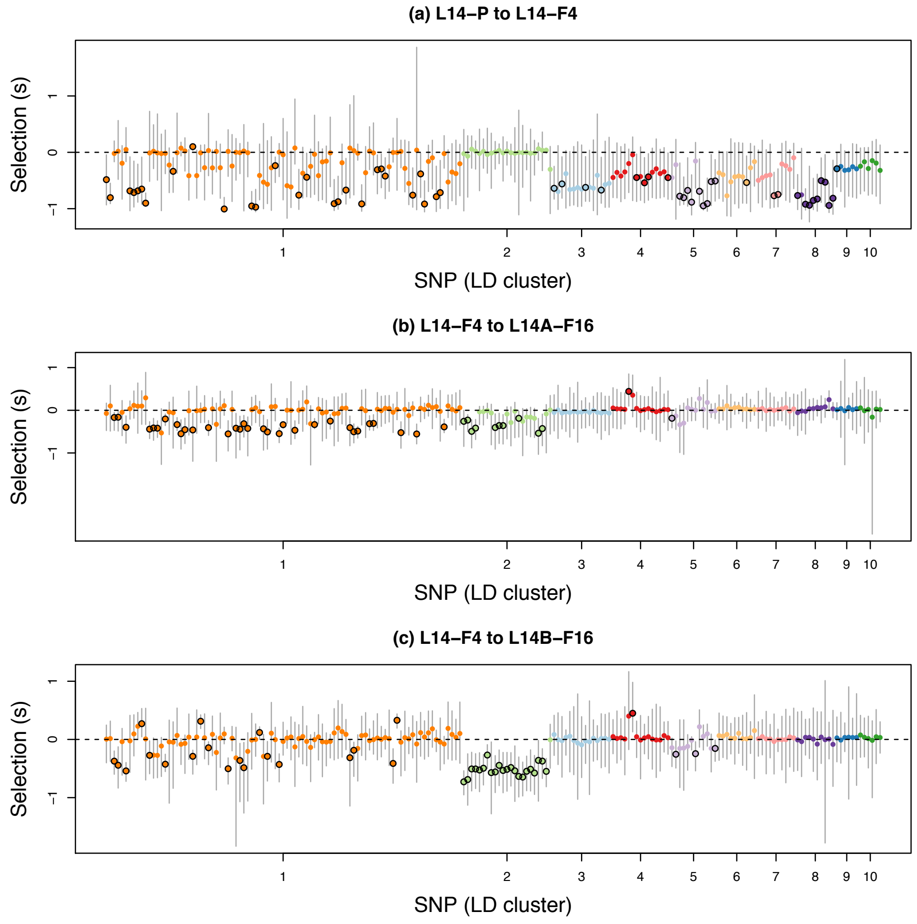


Figure S18: Scatter plots show Bayesian estimates of selection coefficients for the 198 focal SNPs in different generations and sublines with an **unconstrained model** with *a priori* independent estimates of $s$ across sublines and time intervals. Dots and vertical bars denote posterior medians and 95% equal-tail probability intervals (ETPIs), respectively. Colors and the order of SNPs reflect LD cluster membership in the F1 generation. Black circles around dots denote cases where the 95% ETPIs exclude 0. For the purpose of visualization, have polarized estimates of $s$ such that negative values indicate selection favoring the minor allele.

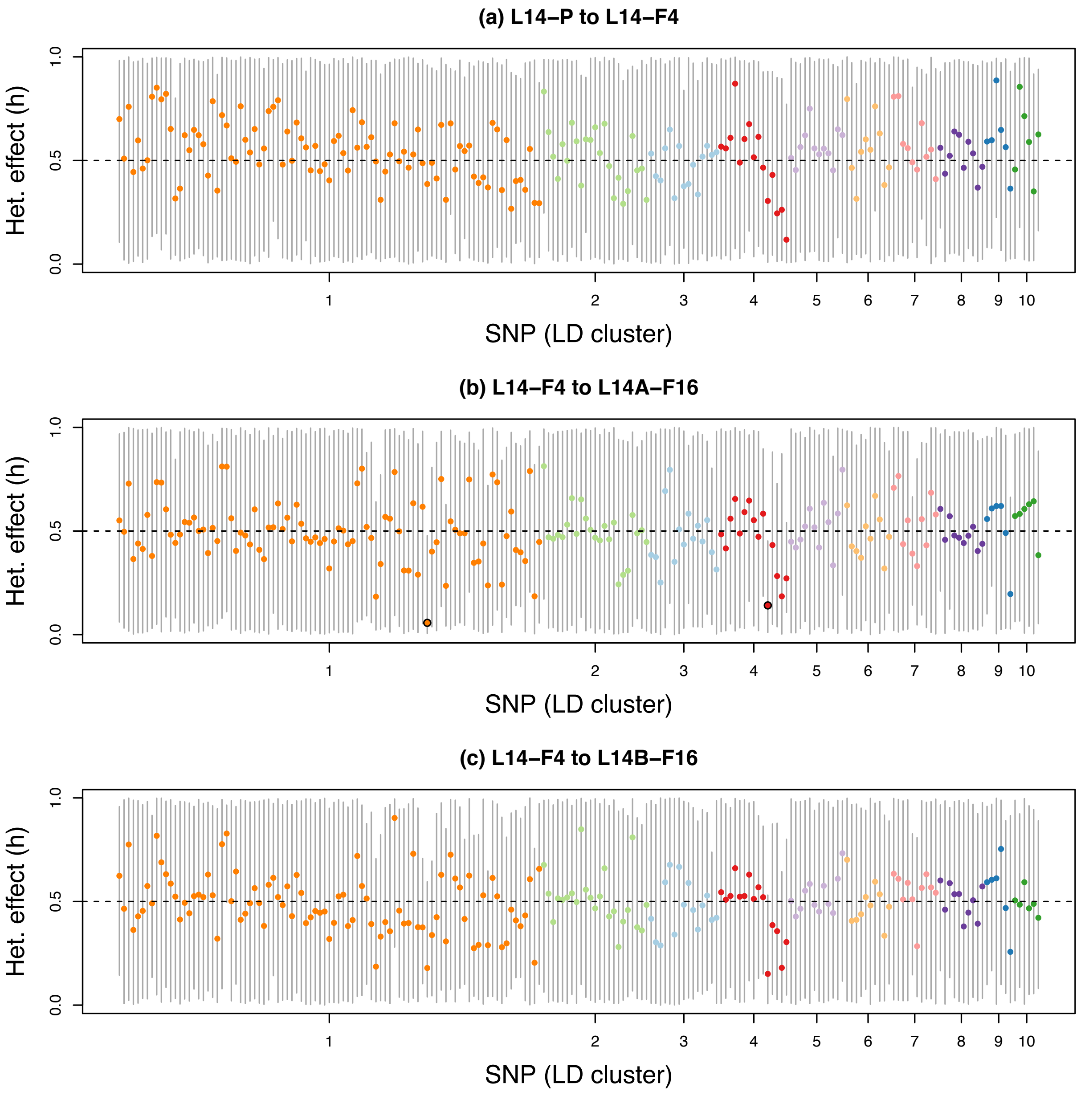


Figure S19: Scatter plots show Bayesian estimates of the heterozygous/dominance effect ($h$) for the 198 focal SNPs in different generations and sublines. Dots and vertical bars denote posterior medians and 95% equal-tail probability intervals (ETPIs), respectively. Colors and the order of SNPs reflect LD cluster membership in the F1 generation. Black circles around dots denote cases where the 95% ETPIs exclude 0.5.

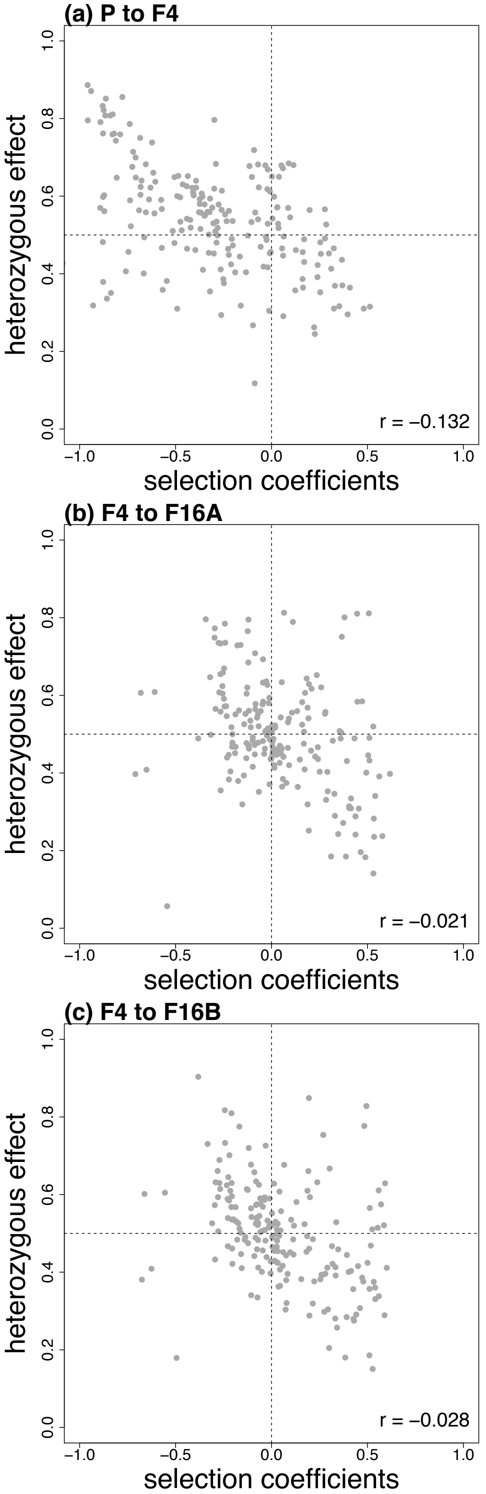


Figure S20: Scatter plots show the relationships between selection coefficient estimates and heterozygous effect estimates for the 198 focal SNPs in different time intervals and sublines. Pearson correlations account for uncertainty in estimates of selection (i.e., they are not based solely on the point estimates shown here).

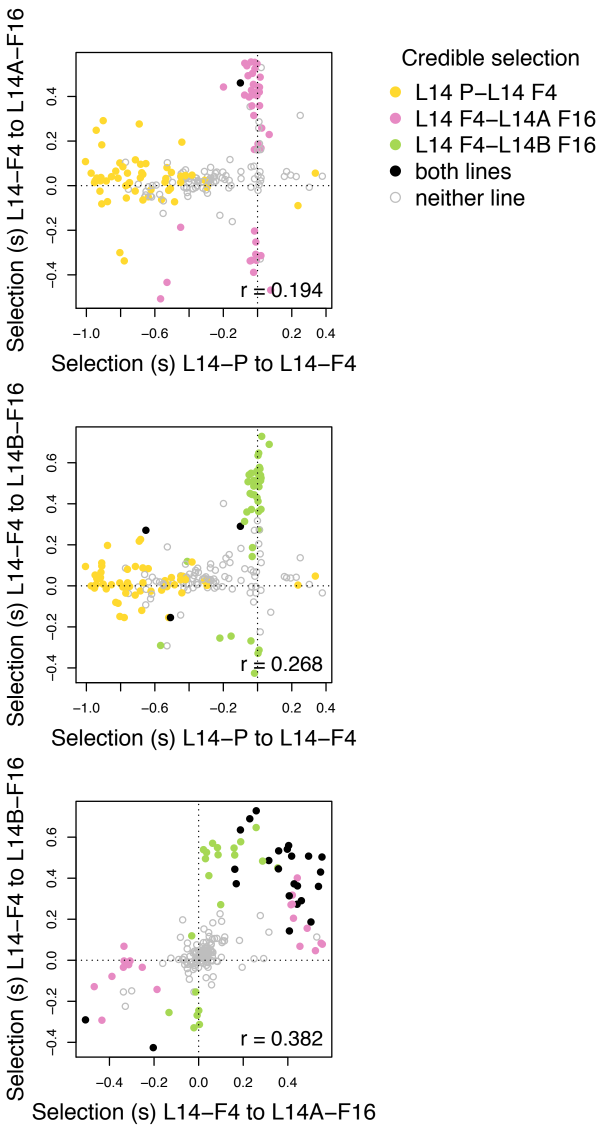


Figure S21: Scatter plots show the relationships between selection coefficient estimates for the 198 focal SNPs in different time intervals and sublines produced by the fully unconstrained ABC simulations. Dots correspond to SNPs and are colored based on whether there was credible evidence of selection in each subline/interval. Pearson correlations account for uncertainty in estimates of selection (i.e., they are not based solely on the point estimates shown here).

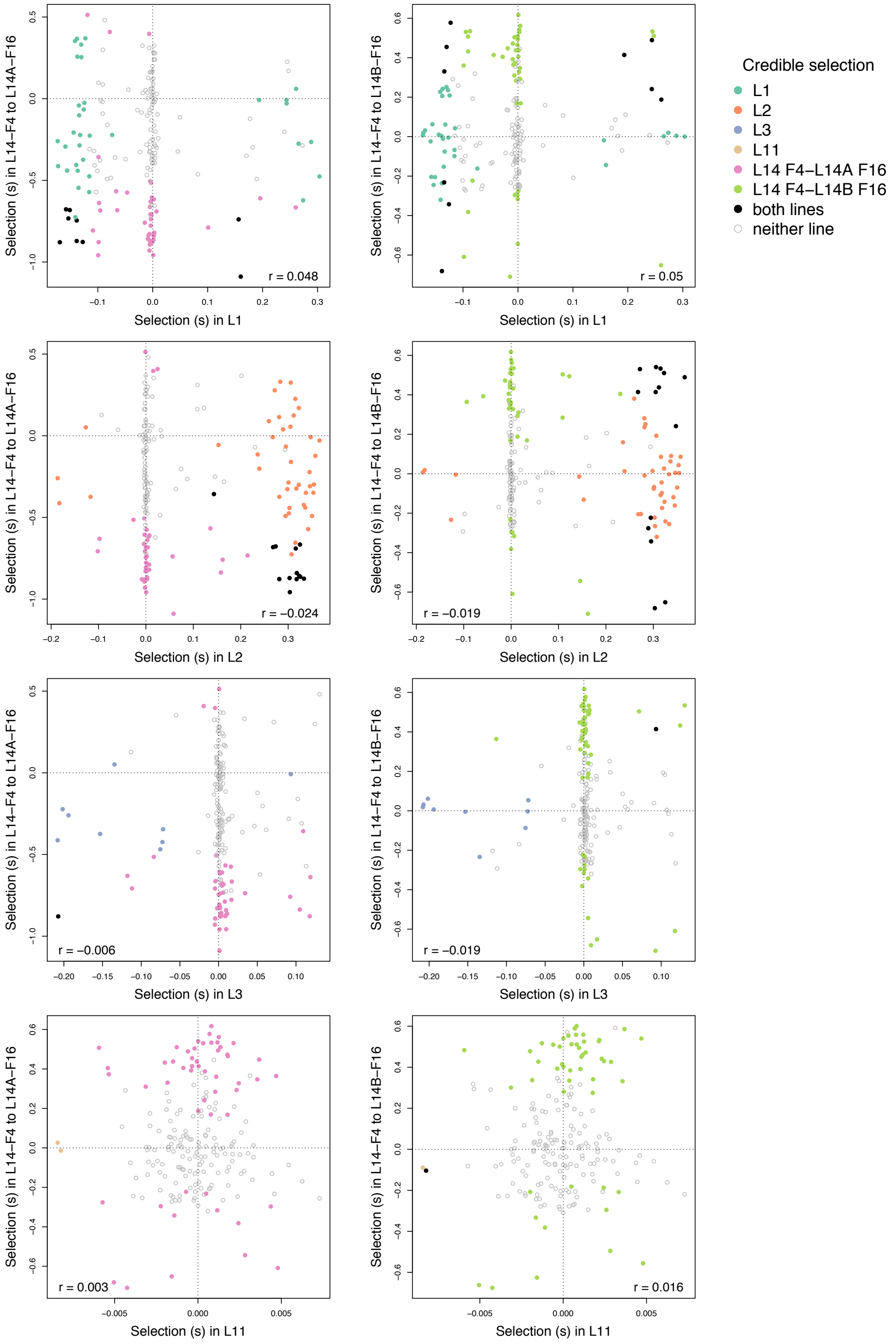


Figure S22: Scatter plots show associations between selection coefficient estimates for the focal SNPs across lines. Results are shown here for all comparisons involving the later stages of rescue in L14 with other lines. For comparisons with lines L1, L2, and L3 the 188 SNPs present in those lines are shown (a single point for L11 was omitted for visualization as it had an extreme but not-credible estimate of $s$). Dots correspond to SNPs and are colored based on whether there was credible evidence of selection in each (sub)line. Pearson correlations account for uncertainty in estimates of selection (i.e., they are not based solely on the point estimates shown here).
